# Dissociation rate compensation mechanism for budding yeast pioneer transcription factors

**DOI:** 10.1101/441469

**Authors:** Benjamin T. Donovan, Hengye Chen, Caroline Jipa, Lu Bai, Michael G. Poirier

## Abstract

Nucleosomes restrict the occupancy of most transcription factors (TF) by reducing binding and accelerating dissociation, while a small group of TFs have high affinities to nucleosome-embedded sites and facilitate nucleosome displacement. To mechanistically understand this process, we investigated two *S. cerevisiae* TFs, Reb1 and Cbf1. We show these factors bind their sites within nucleosomes with similar affinities to naked DNA, trapping a partially unwrapped nucleosome without histone eviction. Both the binding and dissociation rates of Reb1 and Cbf1 are significantly slower at the nucleosomal sites relative to DNA, demonstrating that the high affinities are achieved by increasing the dwell time on nucleosomes to compensate for reduced binding. Reb1 also shows slow migration rate in the yeast nuclei. These properties are similar to human pioneer factors (PFs), suggesting the mechanism of nucleosome targeting is conserved from yeast to human.

---

The fundamental unit of chromatin is the nucleosome, ~147 base pairs (bp) of DNA wrapped around a core of 8 histone proteins ^1^. Extensive contacts between nucleosomal DNA and the histone octamer suppress access to DNA binding proteins, including many transcription factors (TFs) ^2^. To overcome this steric occlusion, TFs take advantage of dynamic nucleosome structural fluctuations, which transiently exposes DNA binding sites for TF binding. However, this site exposure mechanism ^3,4^ results in reduced occupancy relative to naked DNA, since the TF can bind only when the site is partially unwrapped. In addition, it was recently shown that nucleosomes increase TF dissociation rates by orders of magnitude ^5^. In combination, the decreased binding and increased dissociation rates can result in over a 1000-fold reduction in the apparent dissociation constant to nucleosome substrates. For example, Gal4 binds to its DNA target site at picomolar concentrations ^6^ while it requires nanomolar concentrations to bind nucleosomal DNA ^5^. In contrast to TFs like Gal4, pioneer transcription factors (PFs) access their binding sites within nucleosomes as efficiently as fully exposed DNA without the aid of additional factors ^7^. This property is thought to allow PFs to target closed chromatin and prime transcription activation ^8^.

In budding yeast, chromatin is mainly opened by a few highly expressed TFs that can access their nucleosome-embedded binding sites in the genome and establish local Nucleosome Depleted Regions (NDRs) ^9^. How these TFs gain access to their DNA target sites and facilitate nucleosome displacement is not well understood. Two nonexclusive mechanisms may be used during this process. With a ‘passive’ mechanism, TFs can occupy naked DNA when the nucleosome structure is temporarily disrupted by another cellular event (e.g. DNA replication). Note that this mechanism does not require TFs to interact with nucleosomes. Alternatively, TFs may directly bind and invade into nucleosomes. The key to differentiate between these two models is to determine if these TFs, like PFs, can stably engage a nucleosomal template containing their recognition sites ^10^.

One well-studied nucleosome depleting TF is Reb1, a factor essential for yeast viability. Consistent with its ability to displace nucleosomes, most of the Reb1 binding sites in the genome reside in NDRs ^11^. However, ~20% (154/903) of Reb1 binding sites exist within well-positioned nucleosomes and almost half (71/154) of these sites are occupied by Reb1. Reb1 tends to bind the nucleosome near the entry-exit site and was shown to increase the local DNA accessibility ^12^. Overall, these observations suggest that Reb1 may gain access to nucleosomes near the entry-exit site via the site exposure model ^3,12^, but the stability and the kinetics of this interaction are unknown.

In this study, we used a combination of *in vitro* techniques including gel electromobility shift assays (EMSA), ensemble fluorescence, and single-molecule fluorescence to determine if and how Reb1 invades nucleosomes. We find Reb1 accesses its site in both DNA and nucleosomes with similar affinities and targets entry-exit sites by trapping the nucleosome in a partially unwrapped state without evicting histones. Similar to other TFs, the nucleosome site exposure lowers the Reb1 association rate when binding to nucleosome-embedded sites. However, once bound, Reb1 compensates for the reduced association rate with an equally reduced dissociation rate. These properties may be general among these nucleosome-displacing factors, as we show that another *S. cerevisiae* TF, Cbf1, binds nucleosome with similar dynamics. Finally, *in vivo* Fluorescence Recovery After Photobleaching (FRAP) experiments indicate Reb1 undergoes markedly slower exchange within the nucleus relative to other chromatin-interacting proteins. These properties were previously reported for the human PFs ^13,14^. We therefore propose that Reb1 and Cbf1 can function as PFs using this dissociation rate compensation mechanism to efficiently target nucleosomes, partially unwrap nucleosomes, and facilitate the recruitment of transcription regulatory complexes to define NDRs and activate transcription.

## Results

### Reb1 binds DNA and nucleosomes with similar affinities

A defining property of PFs is that nucleosomes do not impede their binding. To determine if Reb1 exhibits this characteristic, we quantified Reb1 affinities to both nucleosome and DNA substrates. For DNA binding experiments, we tested binding to 25 bp oligos containing the Reb1 binding motif. With reconstituted, sucrose gradient purified nucleosomes (**Figure S1**), we tested binding to entry-exit sites because this is where Reb1 preferentially binds *in vivo* ^15^. We tested binding at 4 separate sites positioned in increments of 5 bp throughout the entry-exit region (**Figure 1a**). We refer to these templates as “P*x*,” where “*x*” defines the beginning of the Reb1 binding site in the 601 nucleosome positioning sequence (NPS) (i.e. P3 = binding site starts at 3 bp into the nucleosome). Binding to both DNA and nucleosomes were detected via EMSA, where we titrate Reb1 and observe formation of a slow mobility Reb1 complex (**Figure 1b-c**). For Reb1 binding to nucleosomes, we imaged EMSAs with Cy5-H2A(K119C) and Cy3-DNA fluorescence (**Figure 1c**, **S2**), confirming Reb1 is in complex with nucleosomes. Affinity is measured for each binding reaction by determining the S_1/2_, the concentration at which 50% of the DNA or nucleosomes are bound by Reb1. To DNA, we measured S_1/2 Reb1-DNA +site ESMA_ = 2.3 ± 0.2 nM, while for the 4 nucleosome constructs, we measured: S_1/2 Reb1-Nuc P3 EMSA_= 4.6 ± 0.1 nM, S_1/2 Reb1-Nuc P8 EMSA_= 1.5 ± 0.1 nM, S_1/2 Reb1-Nuc P13 EMSA_ = 8.5 ± 0.2 nM, and S_1/2 Reb1-Nuc P18 EMSA_ = 11.2 ± 0.3 nM. Additionally, we performed EMSAs with DNA and nucleosomes lacking the specific binding site and measured ~10-fold lower affinity to these sequences [S_1/2 Reb1-DNA-site EMSA_ = 21.7 ± 2.3 nM and S_1/2 Reb1-Nuc p- EMSA_ =32.7 ± 0.8 nM (**Figure 1d**, **S3**)]. This property that Reb1 targets DNA and nucleosome with similar affinities mimics PFs in higher eukaryotes ^16^. In contrast, other TFs like Gal4 and LexA, which employ the site exposure model to invade nucleosome entry-exit sites, require over 1,000-fold higher TF concentrations to bind relative to naked DNA ^5^.

**Figure 1:**
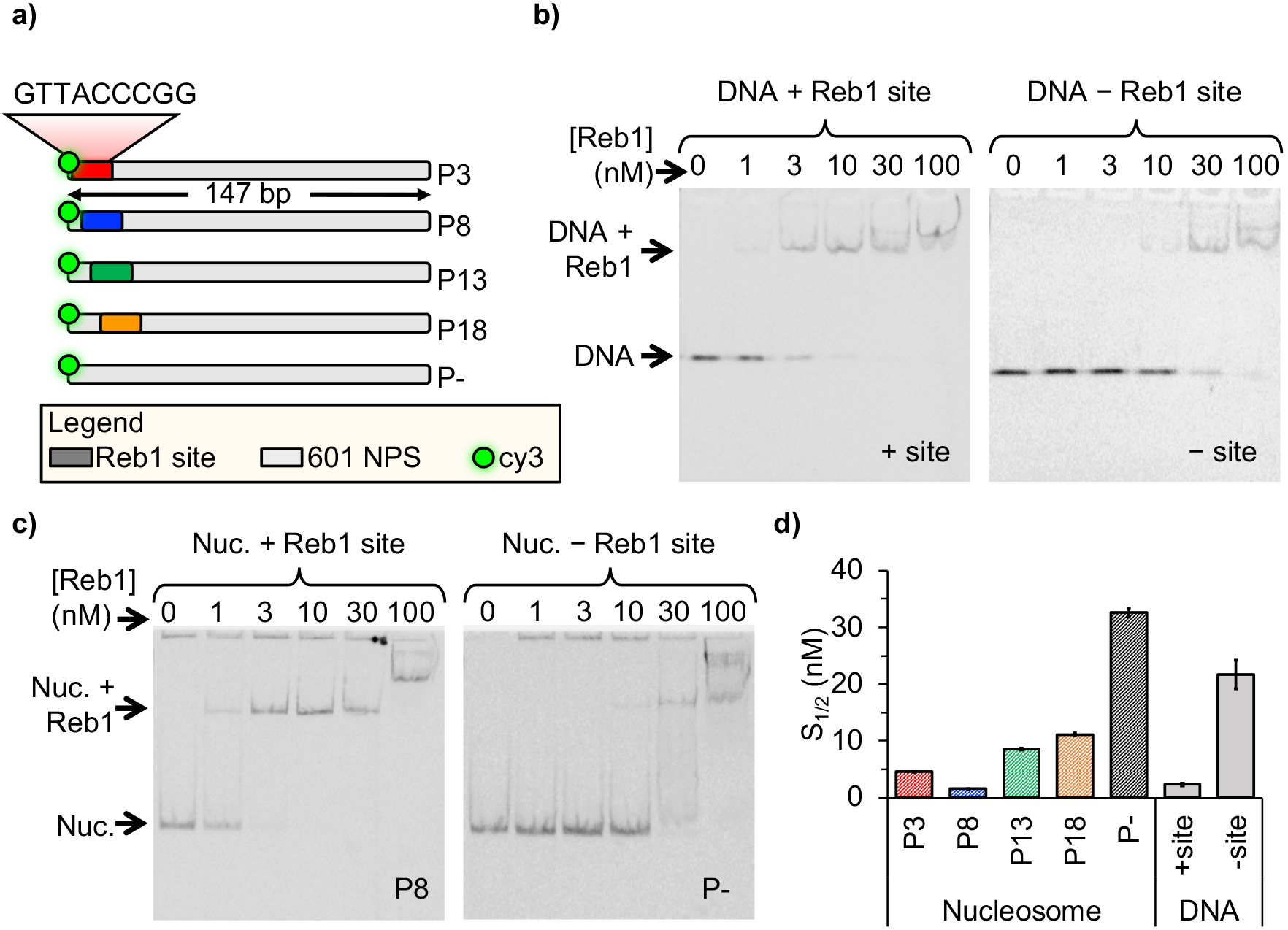
Reb1 binds DNA and nucleosomes with similar affinities. (**a**) Design of the modified “601” nucleosome positioning sequences (NPS) used in this study. Colored rectangles represent the Reb1 binding site at positions P3 (red), P8 (blue), P13 (green) and P18 (gold) within the 601 NPS. The numbers indicates the Reb1 binding site starting position (in number of base pairs into the 601 NPS). (**b**) Cy3 image of the EMSA of Reb1 binding to a 25-bp DNA sequence with (left) or without (right) the Reb1 binding site, (**c**) Cy5 image of the EMSA of Reb1 binding to P8 nucleosomes with (left) or without (right) the Reb1 binding site, (**d**) Quantification of the S_1/2_S determined from the Reb1 EMSAs in B, C and Figure S3 (S_1/2 Reb1-DNA +site EMSA_ = 2.3 ± 0.2 nM, S_1/2 Reb1-DNA - site EMSA_ = 21.7 ± 2.3 nM, S_1/2 Reb1-Nuc P3 EMSA_ = 4.6 ± 0.1 nM, S_1/2 Reb1-Nuc P8 EMSA_ = 1.5 ± 0.1 nM, S_1/2 Reb1-Nuc P13 EMSA_ = 8.5 ± 0.2 nM, S_1/2 Reb1-Nuc P18 EMSA_ = 11.2 ± 0.3 nM, S_1/2 Reb1-Nuc p-EMSA_ =32.7 ± 0.8 nM). These results show that Reb1 binds nucleosomes and DNA site specifically with a similar S_1/2_.

### Reb1 invades nucleosomes by trapping entry-exits sites in a partially unwrapped state

Previous genome-wide studies of nucleosome and Reb1 occupancy suggest that Reb1 gains access to nucleosome entry-exit sites via the site exposure model ^15^. We investigated this binding mechanism through a series of fluorescence resonance energy transfer (FRET) experiments to monitor nucleosomes trapped by Reb1 in partially unwrapped states as previously done for other TFs ^4,17,18^. The donor fluorophore Cy3 was attached to the 5’-end of the DNA adjacent to the Reb1 binding site while the acceptor fluorophore Cy5 was attached to H2A(K119C) (**Figure 2a**). The proximity of these 2 locations within the nucleosome results in high FRET efficiency (85%; **Figure 2b**). For P3 and P8 nucleosomes, titrating Reb1 progressively lowers the FRET efficiency with saturation occurring at ~20%. This relationship fits to a binding isotherm with S_1/2_ values (S_1/2 Reb1-Nuc P3 FRET_ = 7.9 ± 1.3 nM, S_1/2 Reb1-Nuc P8 FRET_ = 2.4 ± 0.3 nM; **Figure 2c**) that agree with S_1/2_ values for the corresponding EMSA measurements (**Figure 1c**). The similarity of the S_1/2_ for Reb1 binding and ΔFRET strongly suggests that Reb1 binding to its target site causes a significant structural change in the nucleosome. In contrast, significantly higher concentrations of Reb1 are required to induce a ΔFRET with P13 nucleosomes (S_1/2 Reb1-Nuc P13 FRET_ = 101.5 ± 19.1 nM), which is ~12-fold higher than the concentration measured by EMSA for Reb1-nucleosome binding (**Figure 1d**). Additionally, we do not observe a significant ΔFRET for P18 nucleosomes. This indicates that Reb1 can bind to sites further into the nucleosomes but does not induce a structural change. Finally, we demonstrate that our observed ΔFRET is site specific, since Reb1 titrations with nucleosomes that do not contain a binding site result in no ΔFRET (**Figure 2b**).

**Figure 2:**
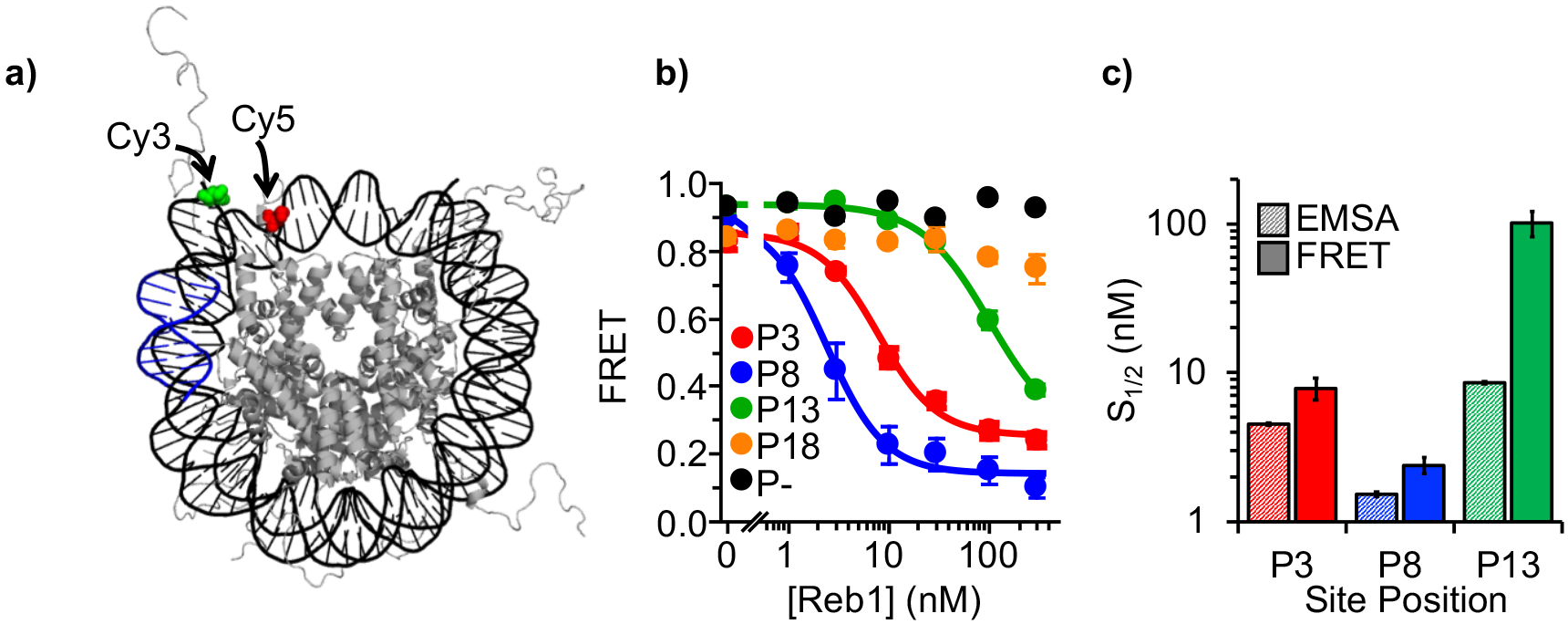
Reb1 binding induces nucleosome structural change. (**a**) Nucleosome structure ^38^ containing the internal FRET pair used in this study. Cy3 is attached to the 5 prime end of the DNA NPS and adjacent to the Reb1 binding site (blue). The octamer is labeled with Cy5 at H2A(K119C). When fully wrapped, the nucleosome is in a high FRET state, (**b**) Nucleosome FRET efficiency measurements while titrating Reb1 with the nucleosome constructs: P3 (red), P8 (blue), P13 (green), P18 (gold), or no binding site (P−, black). Reb1 titrations with P3, P8, and P13 nucleosomes fit to binding isotherms with S_1/2 Reb1-Nuc P3 FRET_ = 7.9 ± 1.3 nM, S_1/2Reb1-Nuc P8 FRET_ = 2.4 ± 0.3 nM, S_1/2 Reb1-Nuc P13 fret_ = 101.5 ± 19.1 nM. We do not observe a significant ΔFRET for P18 and P− nucleosomes. (**c**) Comparison of S_1/2_ values obtained via EMSA and FRET experiments. For P3 and P8 nucleosomes, the FRET S_1/2_ values are in close agreement to the EMSA S_1/2_ values indicating ΔFRET is a measure of Reb1 binding to nucleosomes.

Reb1 occupies nucleosomes at the entry-exit region *in vivo* ^15^. Therefore, we decided to focus on the two Reb1 positions that are closest to the entry-exit region (P3 and P8 nucleosomes) and carried out additional experiments to determine the nature of the Reb1 induce ΔFRET. First, by separately imaging with fluorescence from Cy3-DNA and Cy5-H2A(K119C) in the EMSAs of Reb1-nucleosome binding, we have demonstrated that Reb1 is in complex with nucleosomes. Therefore, the ΔFRET is not due to partial or full nucleosome disassembly (**Figure 1a-b**, **S2, S3**).

It is possible that the observed ΔFRET is due to Reb1 induced a structural change of the H2A C-terminal tail upon binding since the Cy5 fluorophore is positioned in this domain (**Figure S4a**). To test for this, we prepared nucleosomes with a Cy5 fluorophore positioned at H3(V35C). Titrating Reb1 reveals similar nucleosome binding and ΔFRET (**Figure S4b-c**), which rules out the possibility that the ΔFRET is due to a distortion in the H2A C-terminal domain and suggests that the Reb1-induced structural change is between the DNA and the entire octamer.

Another potential explanation for the Reb1-induced structural change is that it traps repositioned nucleosomes. To test this idea, we inserted the Reb1 binding site on the opposite side of the nucleosome from the Cy3 fluorophore and a 20 bp flanking sequence (**Figure S5a**). If Reb1-induced nucleosome structural change is due to octamer translocation, we would detect a decrease in FRET that corresponds with Reb1 binding and repositioning the octamer onto the flanking sequence. While we detect Reb1 binding with EMSA, no ΔFRET is observed (**Figure S5b-c**), which implies that Reb1 binding does not result in repositioned nucleosomes.

These combined results support the conclusion that Reb1 binds to its site within the nucleosome entry-exit region via the site exposure model, where Reb1 traps the nucleosome in a partially unwrapped state. Interestingly, a direct conclusion from the site exposure model is that the Reb1 binding rate should be reduced by the probability the site is exposed, which in this region of the nucleosome is about 100-fold ^17^. Therefore, the site-exposure model alone cannot explain why Reb1 has the same accessibility on partially unwrapped nucleosomes as on naked DNA.

### Reb1 rapidly binds and dissociates at fully exposed DNA binding sites

To investigate how Reb1 can trap a nucleosome in a partially unwrapped state and bind with the same affinity as it binds DNA, we measured Reb1 binding and dissociation kinetics to its site within both DNA and nucleosomes via single molecule Total Internal Reflection Fluorescence (smTIRF) microscopy. Reb1-DNA binding was probed via Protein Induced Fluorescence Enhancement (PIFE), as previously for other TFs ^5,18,19^. Here, the Reb1 binding site was positioned 1 bp away from a Cy3 fluorophore on the 5’ end of the DNA (**Figure 3a**). Titrating Reb1 induces a 1.5-fold increase in Cy3 fluorescence emission, which fits to a binding isotherm with an S_1/2 Reb1-DNA PIFE_ of 5.1 ± 0.2 nM (**Figure 3b**), while without the binding site Cy3 fluorescence does not increase until 100 nM. This agrees with the EMSA binding S_1/2_ (**Figure 3c**), demonstrating that PIFE detects site specific Reb1 binding.

**Figure 3:**
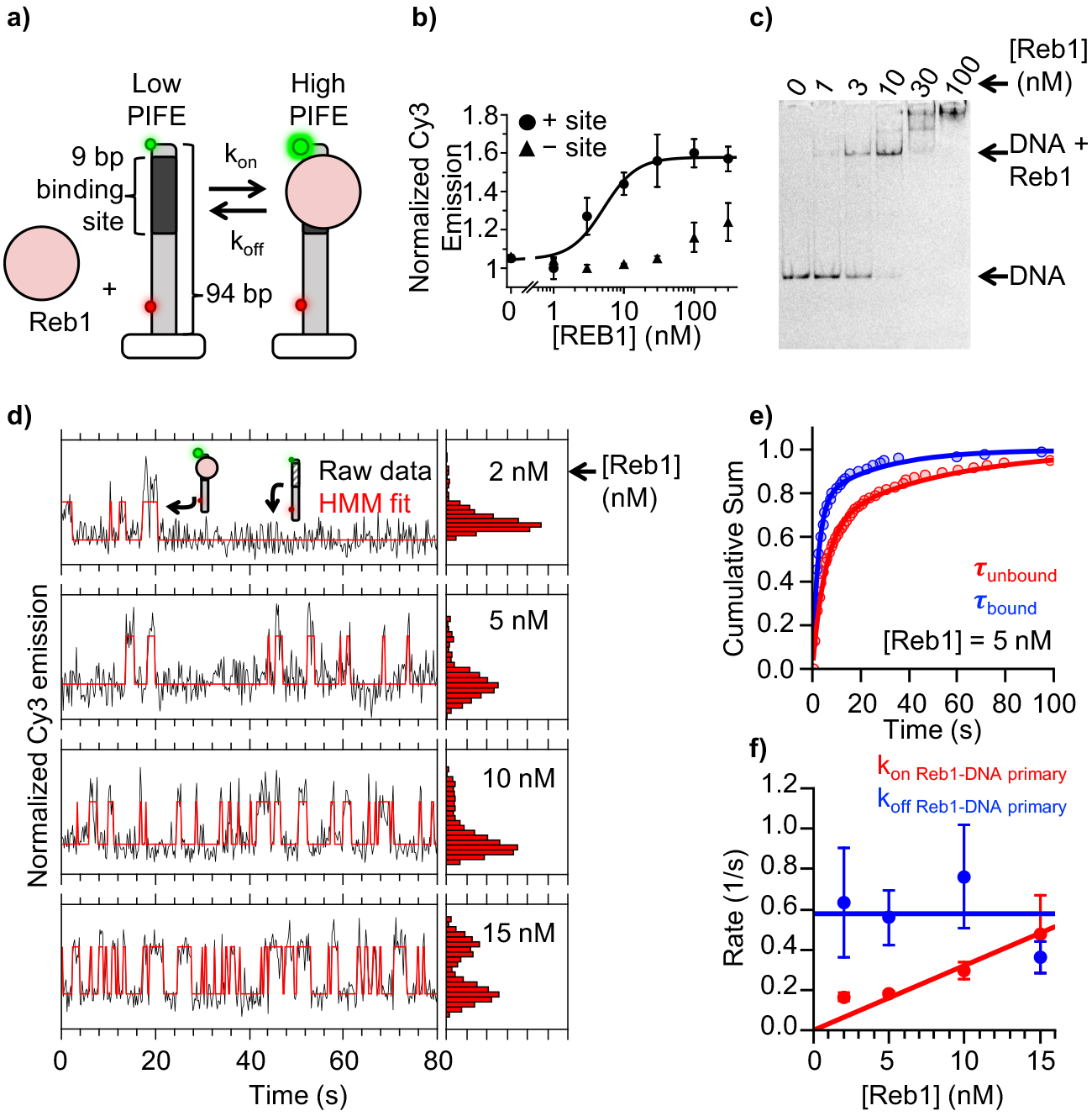
Reb1 rapidly binds to and dissociates from fully exposed DNA binding sites. (**a**) Design of the smPIFE measurements. The 94 bp DNA molecule with the Reb1 binding site 1 bp from the Cy3-labeled 5-prime end was immobilized on a quartz surface through a biotin-streptavidin linkage. DNA molecules are also Cy5 labeled, and we only analyzed molecules with signals in both Cy3 and Cy5. (**b**) Reb1 titration with the smPIFE DNA results in a Cy3 emission increase of ~1.5-fold and fits to a binding isotherm with an S_1/2 Reb1-DNA PIFE_ = 5.1 ± 1.5 nM. Without the binding site, the Cy3 emission does not change until 100 nM, demonstrating the observed PIFE is due to site specific Reb1 binding, (**c**) Cy3 image of the EMSA of Reb1 binding to the smPIFE DNA molecule. Reb1 binds similar to the 25 bp DNA molecule (S_1/2 Rebi-DNA emsa_ = 3.4 ± 0.1 nM). (**d**) Example time traces of single DNA molecules with 4 separate Reb1 concentrations, where the black lines are the Cy3 fluorescence and the red lines are the 2-state Hidden Markov Model fits. As the Reb1 concentration increases, the immobilized DNA molecules shift to the high PIFE state, (**e**) Example cumulative sums of unbound (red) and bound (blue) dwell times that are fit with double exponentials. The Reb1 concentration is 5 nM. (**f**) The primary Reb1-DNA binding (red) and dissociation (blue) rates at 4 Reb1 concentrations. The dissociation rates are constant with an average rate of k_off Reb1-DNA primary_ = 0.58 ± 0.08 s^−1^, while the binding rate increases with Reb1 concentration. The overall binding rate is determined by fitting to a line where the slope represents the binding rate, k_on Rebi-DNA primary_ = 0.032 ± 0.003 s^−1^ nM^−1^.

Next, we performed smTIRF measurements to determine Reb1 binding and dissociation kinetics to and from DNA ^18,20^. The DNA was immobilized on a quartz microscope slide through a biotin-streptavidin linkage and included an 84 bp DNA extension to minimize surface interactions. Additionally, to help ensure the Cy3 signal is due to a surface-tethered DNA molecule, we incorporated an internal Cy5-fluorophore adjacent to the biotin (**Figure 3a**). We only analyzed molecules with both a Cy3 and Cy5 signal.

We measured the time dependent fluorescence from at least 150 single molecules at 4 separate Reb1 concentrations: 2 nM, 5 nM, 10 nM, and 15 nM. These traces fluctuated between a high and low Cy3 fluorescence emission state (**Figure 3d**) and the time spent in the high Cy3 emission state increased with Reb1 concentration. Therefore, we interpreted the high and low Cy3 emission as Reb1 bound and unbound states, respectively, as observed for other TFs^5,18^. We determined the characteristic Reb1 binding and dissociation rates at each Reb1 concentration by compiling the bound and unbound times into separate cumulative sums and fitting these cumulative sums to exponential distributions (**Figure S6a**).

We used the bound time cumulative sums to determine the Reb1 dissociation rate from DNA (**Figure 3e**, **S6a**). Log-likelihood ratio tests indicated the dwell time histograms fit significantly better with double exponential distributions (**Figure S6b**). For all Reb1 concentrations, both rates do not depend on Reb1 concentration (**Figure 3f**, **S7a**), and ~75% of dwell times are associated with the faster rate (**Figure S6b**, **S7b**). This indicates Reb1 mainly remains bound for about a second (k_off Reb1-DNA primary_ = 0.58 ± 0.08 s^−1^; τ_bound_ = 1/koff ≈ 1.7 s), while exhibiting occasional longer binding events (k_off Reb-DNA secondary_ = 0.036 ± 0.005 s^−1^; τ_bound_ ≈ 28 seconds).

To determine Reb1’s binding rate to DNA, we used the unbound dwell time cumulative sum (**Figure 3e**, **S6a**). At each Reb1 concentration, the cumulative sums fit best to a double exponential (**Figure S6b**), where ~75% of unbound times are in the faster population (**Figure S7d**). This primary rate increases with Reb1 concentration (**Figure 3f**), where the slope of the linear fit gives an overall binding rate of k_on Reb1-DNA primary_ = 0.032 ± 0.003 s^−1^ nM^−1^). In contrast, the secondary rate (k_on Reb1-DNA secondary_ = 0.022 ± 0.002 s^−1^) is not Reb1 concentration-dependent (**Figure S7c**), suggesting that it does not represent Reb1 binding. Combined, our results support the conclusion that Reb1-DNA binding and dissociation is highly dynamic on the time scale of ~seconds, and suggests that the primary binding mode of Reb1 is not likely to function as a direct static block of nucleosome occupancy.

### Reb1 binds and dissociates from nucleosomes significantly slower than DNA

To compare Reb1 binding to nucleosomes versus DNA, we next measured Reb1 binding kinetics to nucleosomes using single-molecule FRET ^18^. We focused on the P8 nucleosomes since we could then compare our results to previous investigations of TF-nucleosome interactions ^5, 17,21–23^. The ensemble experiments (**Figure 1**, **2**) demonstrate that Reb1 binding to P8 nucleosomes can be detected by monitoring the change in FRET efficiency. To adapt this experiment for smTIRF, we reconstituted P8 nucleosomes with a DNA that contained an additional 75 bp linker sequence on the side of the nucleosome opposite from the Reb1 binding site and Cy3 fluorophore (**Figure 4a**). With these nucleosomes, we measure an ensemble FRET S_1/2 Reb1-smNuc P8 fret_ of 2.2 ± 0.2 nM, very similar to what was measured for 147 bp P8 nucleosomes (**Figure 4b**) and demonstrate this DNA extension does not impact Reb1 interactions with nucleosomes.

**Figure 4:**
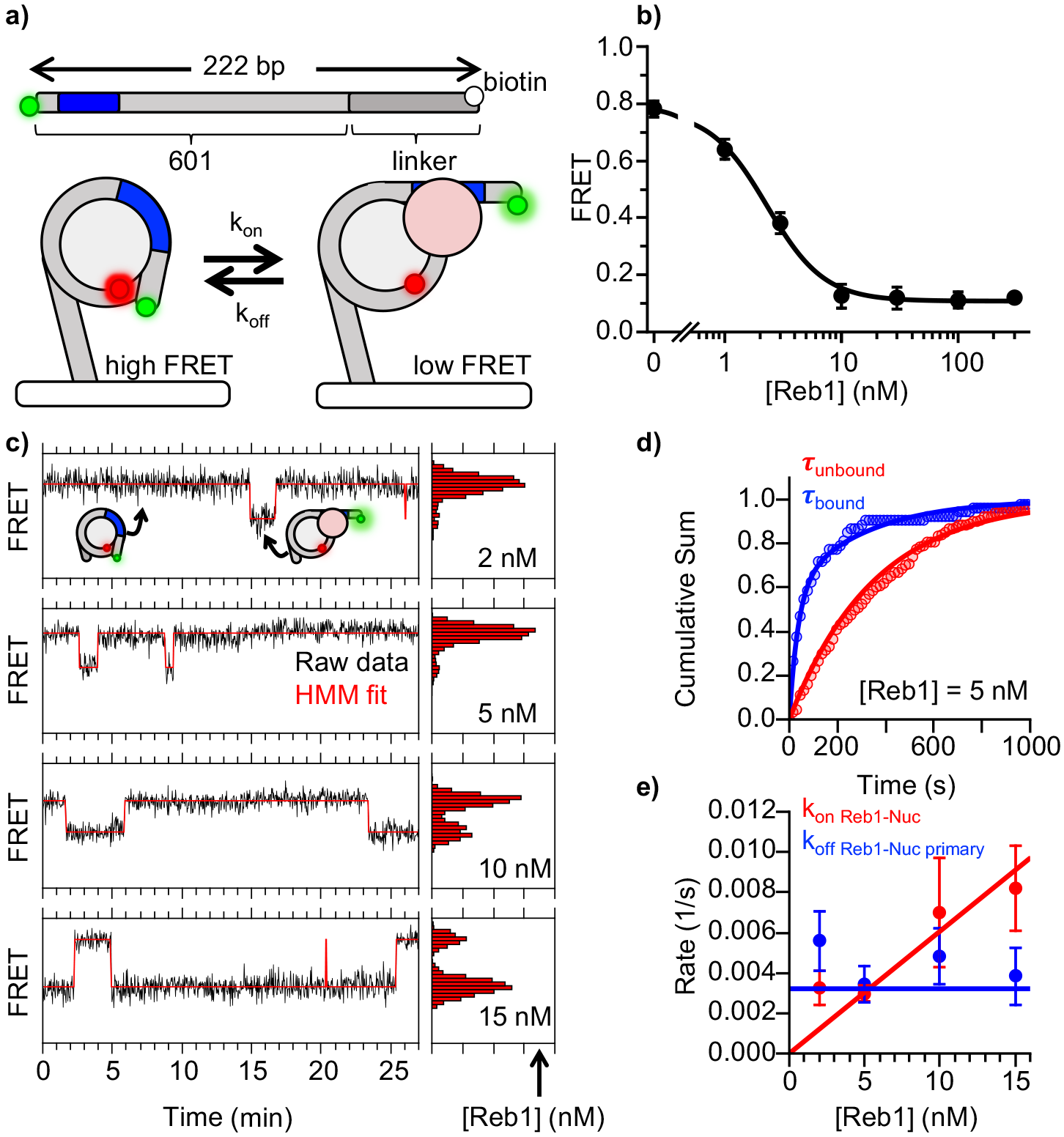
Reb1 binds and dissociates from nucleosomes significantly slower than DNA. (**a**) smFRET P8 nucleosomes are tethered to the microscope surface through an additional 75 bp of DNA extending out of the nucleosome opposite to Cy3 and the Reb1 binding site. The octamer was labeled with Cy5 at H2A(K119C). Reb1 binding traps the nucleosome in a low FRET state, (**b**) Ensemble FRET titration of Reb1 with smFRET nucleosomes. The titration fits to a binding isotherm with an S_1/2 Rebi-smNuc PS FRET_ = 2.2 ± 0.2. This is similar to titrations with nucleosomes containing 147 bp DNA (Figure 2B). (**c**) Example time traces of single nucleosomes with 4 separate Reb1 concentrations, which are fit with a 2-state Hidden-Markov Model. As the Reb1 concentration increases, the immobilized nucleosome shift to the low FRET state, (**d**) Cumulative sums of dwell times in the unbound (red) and states bound (blue), which fit to single and double exponentials, respectively. The Reb1 concentration is 5 nM. (**e**) The primary Reb1 binding (red) and dissociation (blue) rates for increasing Reb1 concentrations. The dissociation rates are constant with an average rate of k_off Rebi-Nuc Primary_= 0.0044 ± 0.0005 s-1, while the binding rates fit to a line with a slope that equals the over binding rate of k_on Reb1-Nuc primary_= 0.0006 ± 0.0001 S^−1^ nM^−1^.

We acquired 30 minute FRET efficiency time series for more than 130 nucleosomes at separate Reb1 concentrations (2, 5, 10, 15 nM). Each nucleosome was immobilized on the microscope slide with a biotin-streptavidin linkage (**Figure 4a**), which fluctuate between high and low FRET states (**Figure 4c**). As we increased the Reb1 concentration, the FRET time series shifts to a larger fraction of time in the low FRET state (**Figure 4c**), indicating Reb1 binds nucleosomes in a partially unwrapped low FRET state. We interpret the high FRET state as a fully wrapped nucleosome without Reb1 bound, and the low FRET state as a Reb1 bound to partially unwrapped nucleosome, as reported for other TFs binding to nucleosomes ^5,18,24^.

The cumulative sums of the unbound (high FRET) dwell times were fit best to single exponential distributions based on log-likelihood ratio tests (**Figure 4d**, **S6a-b**). The rates from the exponential fits increase linearly with increasing Reb1 concentration, where the slope (k_on Reb1-Nuc_ = 0.0006 ± 0.0001 s^−1^ nM^−1^) is the binding rate of Reb1. The Reb1 binding rate is 50-fold lower to its site within the nucleosome relative to DNA, as reported for other TFs binding at this same P8 nucleosome position ^5^. This binding rate reduction confirms that Reb1 binds within nucleosomes by the site exposure mechanism.

The cumulative sums of the bound (low FRET) dwell times were best fit to double exponential distributions based on log-likelihood ratio tests (**Figure 4d**, **S6a-b**). Both rates were independent of Reb1 concentration (**Figure 4e**, **S7c**) and implied Reb1 dissociation rates of k_off Reb1-Nuc primary_ = 0.0044 ± 0.0005 s^−1^ and k_off Reb1-Nuc secondary_ = 0.07 ± 0.02 s^−1^. The majority of bound events (>60%) belonged to the slower population (**Figure S7d**), suggesting that the slower rate is the primary mode for Reb1 dissociation. Interestingly, the Reb1 dissociation rate from nucleosomes is also about 50-fold lower than from DNA. This reduction in dissociation rate compensates for the reduction in binding rate and results in a similar Reb1 binding affinity to its site within nucleosomes and DNA. This result suggests Reb1 interacts with partially unwrapped nucleosomes differently than other TFs like Gal4 and LexA, which exhibit ~1000-fold acceleration in dissociation rates compared to DNA ^5^.

The ratio of the dissociation rate to the binding rate is the apparent dissociation constant, K_D_, which can be compared to the ensemble S_1/2_ measurements. Since binding rates are known to be influenced by restricted diffusion due to surface tethering ^25–27^, we compared the ratio of single molecule apparent K_D_S for binding to nucleosomes and DNA to the ratio of ensemble S_1/2_s for binding nucleosomes and DNA (**Figure S7c**). This will largely remove the impact of the restricted diffusion since the on rates will be impacted similarly for tethered nucleosomes and DNA. When using dominant rates, we find that the relative change in binding affinity between nucleosomes and DNA are in close agreement for single molecule and ensemble experiments. This strongly suggests that k_off Reb1-N uc primary_ is the dominate Reb1 dissociation rate from nucleosomes in solution, and that the surface tethering does not impact the measured dissociation rates.

### Cbf1 also binds and dissociates from nucleosomes significantly slower than DNA

Recent work established that 6 TFs including Reb1 are mainly responsible for NDR generation in *S. cerevisiae* ^9^. To determine if long dissociation rates from binding sites within nucleosomes might be a general property of these factors, we performed binding experiments to DNA and to nucleosomes with Cbf1, another member of this group. Cbf1 exhibits little structural similarity to Reb1; it contains only one DNA binding domain (myc-like), is significantly smaller (39 kDa), and binds to DNA as a dimer ^28^. However, similar to Reb1, its N-terminus is negatively charged and predicted to be unstructured^29^.

EMSA and ensemble PIFE measurements indicate Cbf1 tightly binds DNA (S_1/2 Cbf1-DNA PIFE_= 1.3 ± 0.3 nM) and that binding can be detected through Cy3 PIFE (**Figure 5a**, **S8a**). We then performed smPIFE experiments and detected fluctuations representative of binding as we did with Reb1 (**Figure 5c**). Similarly to Reb1, dwell time cumulative sums were fit to double exponential distributions (**Figure 5d**, **S8c-f**), suggesting that Cbf1 primarily undergoes short binding events (k_off Cbf1-DNA primary_= 0.30 ± 0.05 s^−1^) and exhibits occasional, long dwell times (k_off Cbf1-DNA secondary_ = 0.034 ± 0.004 s^−1^). Additionally, we detect two separate binding rates (k_on Cbf1-DNA primary_ = 0.025 ± 0.006 s^−1^ nM^−1^, k_on Cbf1-DNA secondary_= 0.024 ± 0.003 s^−1^) where only the primary rate depends on Cbf1 concentration.

**Figure 5:**
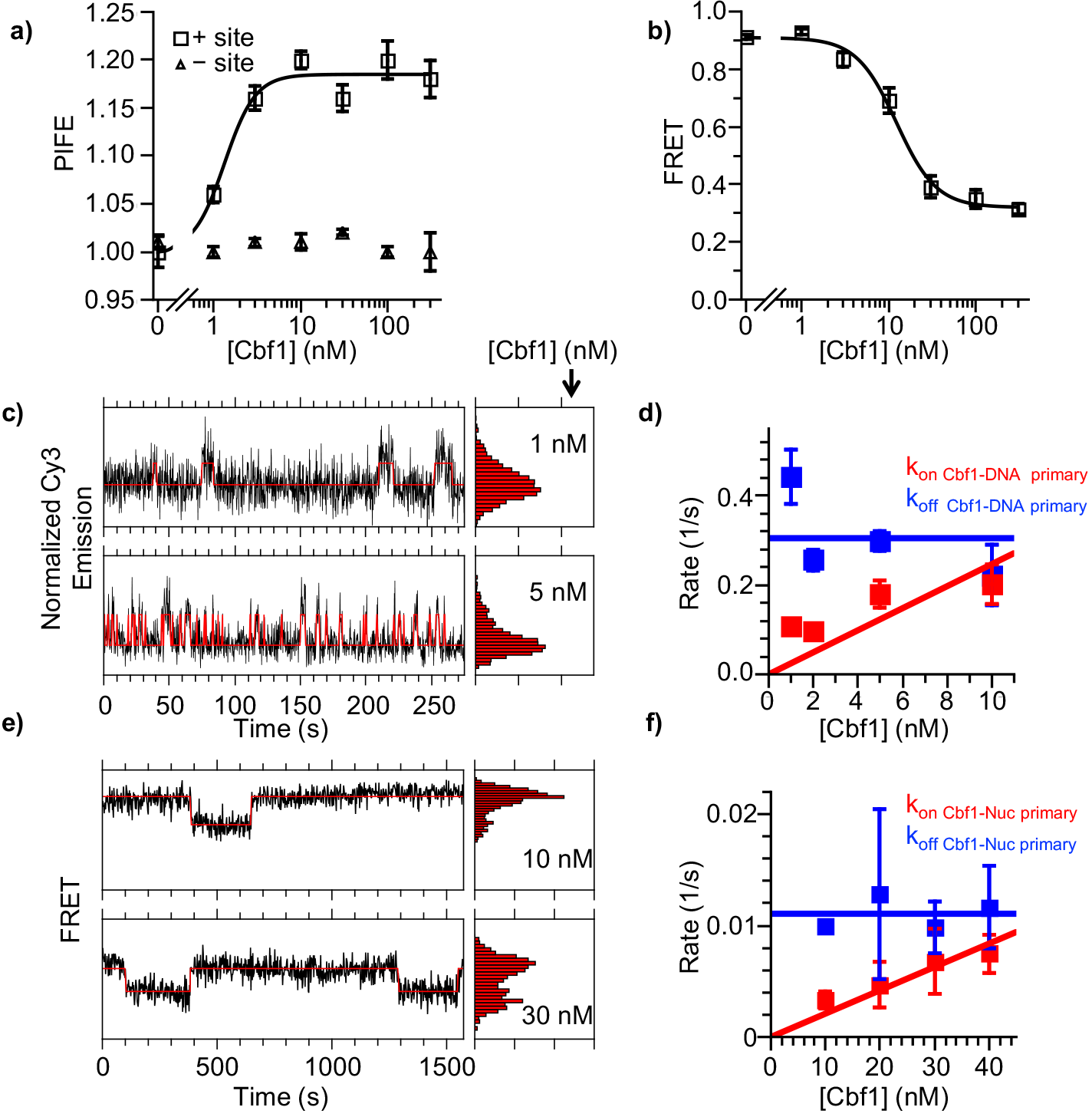
Cbf1 also binds and dissociates from nucleosomes significantly slower than DNA. (**a**) Ensemble PIFE measurement of a Cbf1 titration with a 94 bp DNA with and without the Cbf1 binding sites 1 bp from the 5 prime end cy3 labeled. The normalized PIFE fits to a binding isotherm with an S_1/2 cbfi-DNA PIFE_ = 1.3 ± 0.3 nM. Without the binding site, the Cy3 emission does not change demonstrating the observed PIFE is due to site specific Cbf1 binding, (b) Cbf1 titration with Cy3-Cy5 labeled nucleosomes with the Cbf1 sites at P8. The FRET fits to a binding isotherm with an S_1/2 cbfi-smNuc FRET_ = 12.3 ± 1.6 nM. (**c**) Example time traces of single DNA molecules with 2 separate Cbf1 concentrations, where the black lines are the Cy3 fluorescence and the red lines are the 2-state Hidden Markov Model fits. As the Cbf1 concentration increases, the immobilized DNA molecules shift to the high PIFE state, (**d**) The Cbf1-DNA primary binding and dissociation rates for increasing concentrations of Cbf1. These were determined from cumulative sums of Reb-DNA bound and unbound dwell times that were fit to double exponentials. The primary dissociation kinetics (blue) were constant with an average of k_off cbfi-DNA primary_ = 0.30 ± 0.05 s^−1^, while the primary binding kinetics (red) fit to a line with a slope that equals the overall binding rate of k_on Cbf1-DNA primary_ = 0.025 ± 0.006 s^−1^ nM^−1^. (**e**) Example time traces of single nucleosomes with 2 separate Cbf1 concentrations, where the black lines are the FRET efficiency data and the red lines are the 2-state Hidden Markov Model fits. As the Cbf1 concentration increases, the immobilized nucleosome shift to the low FRET state, (**f**) The Cbf1-nucleosome binding and dissociation rates for increasing concentrations of Cbf1. These were determined from cumulative sums of Reb1-nucleosome bound and unbound dwell times that were fit to single exponentials. The dissociation kinetics (blue) were constant with an average of k_off cbfi-Nuc_ = 0.0111 ± 0.0007 s^−1^, while the binding kinetics (red) fit to a line with a slope that equals the overall binding rate of k_on cbfi-Nuc_ = 0.00021 ± 0.00002 s^−1^ nM^−1^.

We then used both EMSA and ΔFRET to detect Cbf1 binding to Cy3-Cy5 labeled nucleosomes with the Cbf1 binding site at the P8 position. We used the same Cy3 and Cy5 label positions as in the Reb1 FRET measurements (**Figure 2a**). Titrating Cbf1 resulted in a significant reduction in FRET with an S_1/2 Cbf1-Nuc fret_ = 12.3 ± 1.6 nM, which is similar to the EMSA measurements and indicates that FRET can be used to measure Cbf1 binding (**Figure 5b**, **S8b**). Comparison of these ensemble measurements reveal a ~10-fold lower binding affinity of Cbf1 to P8 nucleosomes as compared to naked DNA. We then used smFRET to determine the kinetic rates of Cbf1 binding to and dissociating from P8 nucleosomes (**Figure 5e**). The cumulative sums were fit best with single exponential distribution (k_on Cbf1-Nuc_ = 0.00021 ± 0.00002 s^−1^ nM^−1^ and k_off Cbf1-Nuc_= 0.0111 ± 0.0007s^−1^) (**Figure 5f**, **S8c-d**). As with Reb1, using the dominant rates from these measurements, we determined that the relative change in Cbf1 binding affinity to DNA and nucleosomes is consistent between single molecule and ensemble measurements (**Figure S8g**).

Similar to Reb1, we detect a ~120-fold slower Cbf1 binding rate to nucleosomes as compared to DNA, which indicates Cbf1 also gains access to nucleosomes via the site exposure mechanism. Interestingly, the Cbf1 primary dissociation rate is ~25-fold lower from nucleosomes relative to DNA, which is qualitatively similar to Reb1 and in stark contrast to the orders of magnitude increase in dissociation rate of the Gal4 and LexA TFs ^5^. The decreased rate of Cbf1-nucleosome dissociation partially compensates for the decreased binding rate and explains why Cbf1 binds its site within nucleosomes only about 10-fold weaker than DNA as compared to the orders of magnitude decrease in occupancy for both Gal4 and LexA ^3,5,6^. Combined, these results on Cbf1 and Reb1 indicate that, similar to PFs in higher eukaryotes, they have high affinity to nucleosomal substrates, and that they achieve high affinity by reducing their dissociation rates to compensate for their reduced binding rates.

### Reb1 slowly exchanges *in vivo*

To further investigate our observation that Reb1 can function as a PF, we carried out FRAP measurements. Previous FRAP measurements of mammalian GFP tagged PFs show that they exchange with a characteristic recovery time significantly slower than other TFs in the nuclei ^13,30^. We carried out FRAP measurements of endogenously expressed GFP tagged Reb1 in *S. cerevisiae*, and observed that Reb1 fluorescence recovers with a half life of 25.8 ± 2.5 seconds (**Figure 6**), which is similar to the exchange times observed for the mammalian PF, FoxA. We could not get high-quality FRAP data on Cbf1, which is less abundant than Reb1.

**Figure 6:**
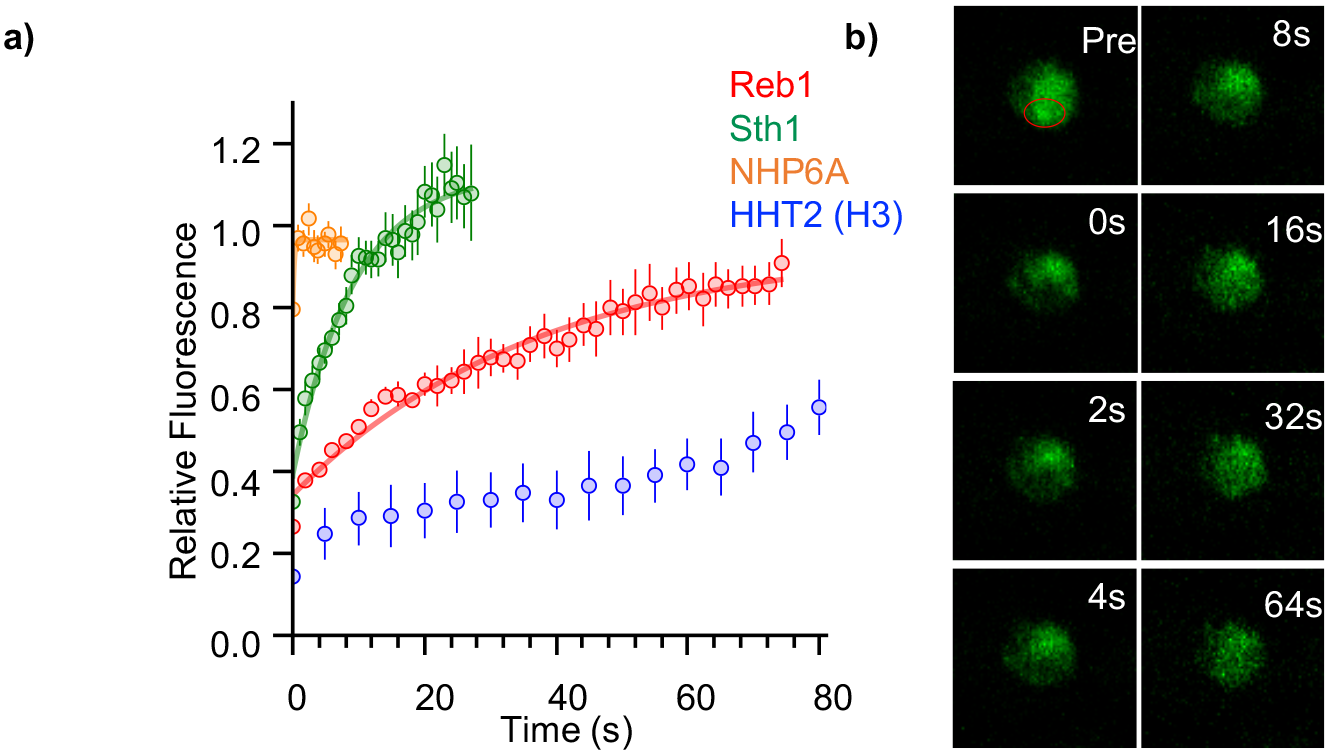
Reb1 slowly exchanges in vivo. (**a**) Recovery curves of Reb1 (red), HHT2 (Blue), Sth1 (green), and NHP6A (gold) after photobleaching. Reb1 **τ**2 = 25.8 ± 2.5 s, Sth 1 **τ**_1/2_ = 7.8 ± 0.7 s, NHP6A **τ**_1/2_ = 0.2 ± 0.1 s. (**b**) Fluorescence images of GFP-labeled Reb1 during a FRAP experiment. The bleached region is indicated with a red circle.

For comparison, we performed FRAP measurements of three additional endogenously expressed GFP tagged proteins that interact with chromatin: histone H3, Sth1, and Nhp6A. These factors were chosen because they are abundant nuclei proteins, and are expected to have different levels of chromosome engagement. H3 is stably integrated into chromatin and has been previously reported to exchange on the hour time scale in mammalian cells ^31^. Sth 1 is a subunit of the nucleosome remodeling complex RSC, which has strong nucleosome interactions, but interacts with chromatin transiently ^32,33^.

Nhp6A is a high-mobility group protein that binds to DNA with low sequence-specificity^34^, and its mammalian homolog, HMGB1, was shown to have rapid second FRAP recovery rates ^13^. We find that the FRAP half-time of H3 to be much longer than the minute time scale, the length of our experiment. In contrast, the half-lives for Sth1 and Nhp6A are 7.8 and <1s respectively (**Figure 6**, **S9**). This shows Reb1 exchanges faster than the chromatin forming protein, histone H3, but slower than other transcription regulatory proteins that transiently interact with chromatin. This result provides additional evidence that Reb1 functions *in vivo* similarly to mammalian PFs, where it exchanges slowly compared to other transcription regulatory complexes.

## Discussion

Here we combine ensemble, single molecule and live cell fluorescence studies to investigate mechanistically how the budding yeast TFs, Reb1 and Cbf1, interact with nucleosomal templates. We find that, similar to PFs, Reb1 and Cbf1 occupy sites within the nucleosome with similar affinities as to naked DNA. These factors invade the nucleosome and trap it in a partially unwrapped state via the site exposure model, which results in a 100-fold or higher reduction in the binding rate ^17,35^. Interestingly, Reb1 completely and Cbf1 partially compensate for this *binding* rate reduction by reducing their *dissociation* rates (**Figure 7a**). This dissociation rate compensation mechanism explains how a TF can both have similar affinity on naked vs. nucleosomal DNA (**Figure 7c**), as has been proposed for the human PF FoxA ^14^. Although the yeast and human PFs have similar properties on nucleosome binding, it should be noted that their impact on nucleosome structure and dynamics appears to be distinct. FoxA appears to trap mobile nucleosomes into distinct positions along the DNA ^14,36^ and compete off linker histones ^36^, while Oct4 and Sox2 do not appear to disrupt nucleosomes upon binding the DNA entry-exit region ^16^. The dissociation rate compensation mechanism by which PFs efficiency target their DNA sites in nucleosomes appears to be conserved from yeast to human, while their impact on nucleosomes structure and binding is much more diverse.

**Figure 7:**
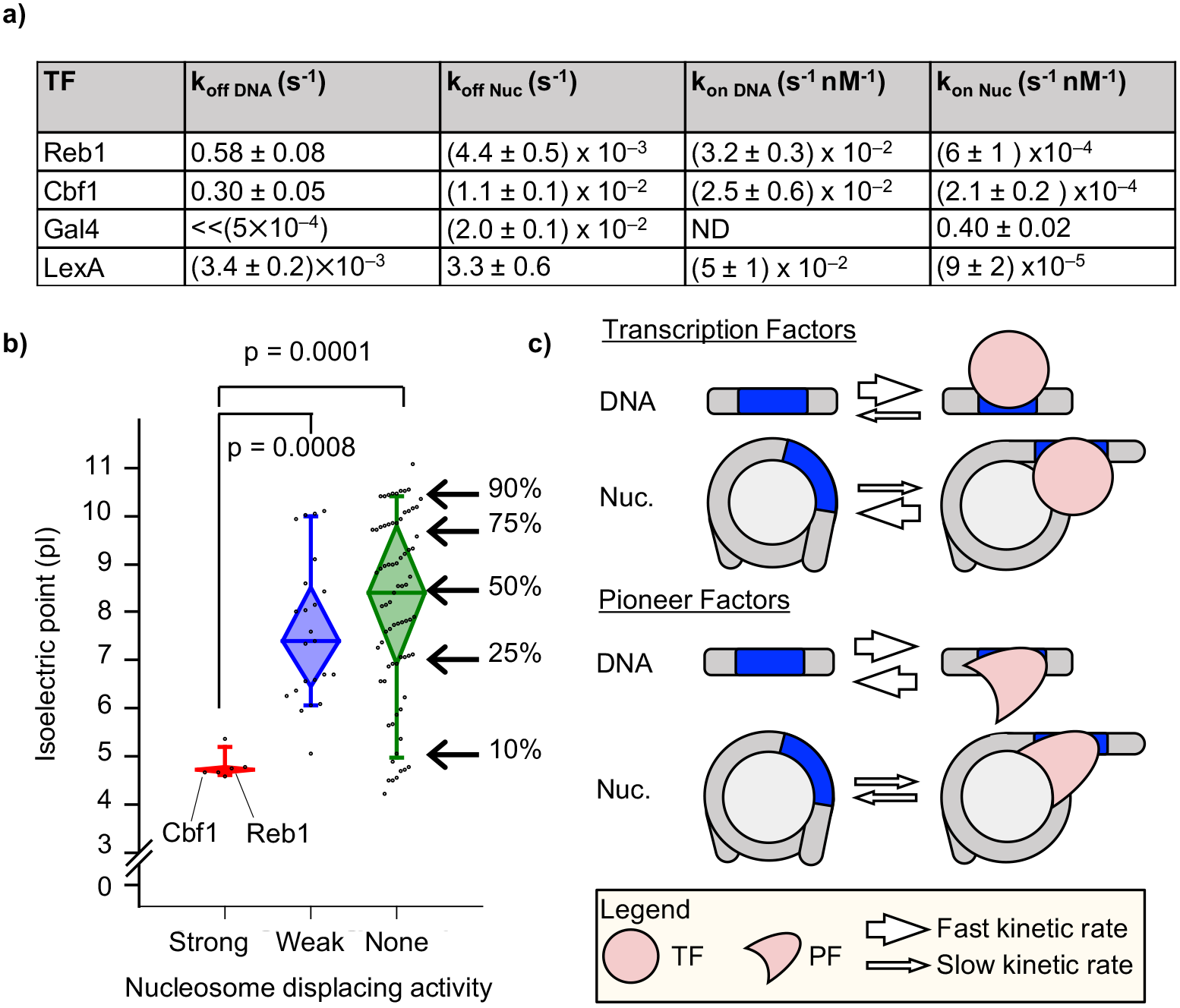
Dissociation rate compensation mechanism for yeast pioneer transcription factors. (**a**) Table of Reb1, Cbf1, Gal4 ^5^ and LexA ^5^ binding and dissociation rates with DNA and P8 nucleosomes. (**b**) The isoelectric point for budding yeast TFs that have strong (red), weak (blue) or no (green) nucleosomes displacing activity ^9^. Reb1 and Cbf1 are part of two of the six with strong nucleosome displacing activity. These factors are significantly more acidic than the groups with weak and no nucleosome displacing activity, (**c**) The dissociation rate compensation mechanism: (top) For traditional TFs, nucleosomes decrease TF binding rates and increase TF dissociation rates, which can reduce the over TF affinity by orders of magnitude, (bottom) For PFs, nucleosomes similarly decrease PF binding rates. However, nucleosomes decrease PF dissociation rates, which compensates for the reduced PF binding rate so the over affinity is similar between nucleosomes and DNA and allows PFs to efficiently trap nucleosomes in partially unwrapped states.

Our results on Reb1 and Cbf1 dissociation rates are strikingly different than previous results on TFs such as Gal4 and LexA, where their dissociation rates are much higher from nucleosomes than DNA. This acceleration has been proposed to be due to both nucleosome rewrapping competing with TF partially bound states and through disfavorable TF-nucleosome interactions such as steric clash ^37^. The reason why Reb1 and Cbf1 dwell much longer on their nucleosomal sites is still not clear. Since the human PF FoxA stabilizes its binding to nucleosomes by contacting H3 via its C-terminus ^36^, we suspect that similar mechanism may also stabilize Reb1 and Cbf1 binding. Interestingly, both Reb1 and Cbf1 are part of a group of 6 budding yeast TFs that can efficiently create NDRs ^9^, and all 6 of these TFs are highly acidic as compared to proteins that do not have such activity (**Figure 7b**). Perhaps, Reb1 and Cbf1 form favorable electrostatic interactions with the basic histone surface that is exposed in a partially unwrapped nucleosomes (**Figure 7c**), which prevents nucleosome rewrapping and significantly reduce their dissociation rates.

The findings above have significant implications on how certain TFs generate NDRs *in vivo*. A previous study proposed that TFs in *S. cerevisiae* can invade into nucleosomes passively by trapping transiently exposed naked DNA during replication or other histone-turnover events ^9^. However, it does not exclude the possibility that some TFs can directly bind and invade into nucleosomes. Our observation that Reb1 and Cbf1 stably engage nucleosome provides support for the latter. Given that these two factors do not evict histones *in vitro* (nor do other PFs), we do not think it is likely that these factors can cause spontaneous nucleosome disassembly *in vivo*. Instead, the long dwell time of these factors on nucleosomes may allow time to recruit other factors, such as histone chaperones or nucleosome remodelers, to establish NDRs. Future *in vivo* and *in vitro* measurements are needed to investigate this further.

## Acknowledgements

We thank the members of the Poirier and Bai Labs for help for helpful discussions. This work was supported by the NIH grants R01 GM121858 (to LB and MGP), R01 GM121966 (to MGP), T32 GM086252 (to BTD) and the NSF grant 1516979 (to MGP).

## Author Contributions

BTD, MGP and LB designed the study. BTD, MGP, LB wrote the paper. BTD performed and analyzed all gel shift, ensemble fluorescence and single molecule experiments. CJ provided support for all of the fluorescence and gel shift experiments. HC prepared, carried out and analyzed all of the FRAP measurements. All authors reviewed the results and approved the final version of the manuscript.

## Materials and Correspondence

Michael G. Poirier

## Methods

### Preparation of Reb1

Reb1 was cloned into pHIS8 and expressed as previously described ^39^. Briefly, Reb1 was expressed in *Escherichia coli* BL21(DE3) cells (invitrogen) by inducing at OD_600_ = 0.4-0.6 with 1 mM IPTG for 3 hours at 37C. Cells were resuspended in 15 mL lysis buffer (50 mM sodium phosphate monobasic pH 8, 300 mM NaCl, 10 mM imidazole, 1 mM PMSF, 1 mM DTT) per 600 mL culture and lysed by sonication. Cell debris was removed by centrifugation, loaded onto a 5 mL HisTrap HP Ni-NTA column (GE healthcare), and eluted with elution buffer (50 mM sodium phosphate monobasic pH 8, 300 mM NaCl, 250 mM imidazole, 1 mM PMSF, 1 mM DTT). Peak fractions were concentrated and further purified with a superdex s200 10/300 size exclusion column that was equilibrated with the Reb1 storage buffer (20 mM HEPES-NaOH pH 7.5, 350 mM NaCl, .1% Tween-20). Pure fractions, as determined by coomassie SDS PAGE, were pooled, concentrated, glycerol added to final concentration of 10%, flash-frozen, and stored at −80C.

### Preparation of Cbf1

Cbf1 was a gift from S Diekmann and was expressed and purified as previously described ^28,40^. Briefly, Cbf1 was cloned into pET28a and expressed in *Escherichia coli* BL21(DE3) cells (invitrogen) by inducing at OD_600 nm_ = 0.4-0.6 with 0.5 mM IPTG for 4 hours at 37 C. Cells were resuspended in buffer A (50 mM Na_2_HP0_4_ (pH 7.5), 300 mM NaCl, 5 mM imidazole, 10% glycerol, 1 mM PMSF, 20 ug/mL pepstatin, 20 ug/mL leupeptin), lysed by sonication, and cell debris was removed by centrifugation (4 C, 23,000 × G, 20 min). After centrifugation, lysate was loaded onto a 5 mL HisTrap HP Ni-NTA column (GE healthcare) and washed with 40 mL buffer A, 120 mL buffer B (50 mM Na_2_HP0_4_ (pH 7.5), 300 mM NaCl, 60 mM imidazole, 10% glycerol, 1 mM PMSF, 20 ug/mL pepstatin, 20 ug/mL leupeptin), and eluted with with buffer C (50 mM Na_2_HP0_4_ (pH 7.5), 300 mM NaCl, 340 mM imidazole, 10% glycerol, 1 mM PMSF, 20 ug/mL pepstatin, 20 ug/mL leupeptin). Pure fractions (as determined by SDS PAGE) were pooled, and imidazole was removed by washing with Buffer D (50 mM Na_2_HP0_4_ (pH 7.5), 300 mM NaCl, 10% glycerol, 1 mM PMSF, 20 ug/mL pepstatin, 20 ug/mL leupeptin) in a 10 K amicon (Millipore).

### Preparation of DNA molecules

DNA molecules for PIFE, FRET, and EMSA experiments were prepared by PCR with cy3/cy5/biotin-labeled oligonucleotides (Sigma) from a plasmid containing the 601 nucleosome positioning sequence (NPS) with a consensus Reb1 or Cbf1 binding site at various positions. For Reb1 experiments, the potential Reb1 binding site at pos. 87-93 in the 601 was removed by site-directed mutagenesis. Oligonucleotides (**Figure S10**) were labeled with Cy3 or Cy5 NHS ester (GE healthcare) at an amino group attached at the 5’-end or at an amine modified dT, and purified by HPLC with 218TP C18 column (Grace/vydac). Following PCR amplification, DNA molecules were purified using a MonoQ column (GE healthcare).

### Preparation of Histone Octamers

Human recombinant histones were expressed and purified as previously described ^41^. Expression vectors were generous gifts from Dr. Karolin Luger (University of Colorado) and Dr. Jonathan Widom. Mutation H3(C110A) was introduced by site-directed mutagenesis (agilent). The histone octamer was refolded by adding each of the histone together at equal molar ratio and purifying as previously described ^41^. H2A(K119C) and H3(V35C)-containing HO were labeled with Cy5-maleamide (GE Healthcare) as previously described ^42^.

### Preparation of nucleosomes

Nucleosomes were reconstituted from Cy3-labeled DNA and purified Cy5-labeled histone octamer (HO) by double salt dialysis as previously described ^42^. Dialyzed nucleosomes were loaded onto 5-30% sucrose gradients and purified by centrifugation on an Optima L-90 K Ultracentrifuge (Beckman Coulter) with a SW-41 rotor. Sucrose fractions containing nucleosomes were collected, concentrated, and stored in .5× TE pH 8 on ice.

### Electrophoretic mobility shift assays

0.5 nM DNA or nucleosomes were incubated with 0-100 nM Reb1 in 10 mM Tris-HCl pH 8, 130 mM NaCl, 10% glycerol, 0.0075% v/v Tween-20 for at least 5 minutes and then resolved by electrophoretic mobility shift assay (EMSA) with a 5% native polyacrylamide gel in .3× TBE.

### Ensemble PIFE measurements

Reb1 binding to its target site on Cy3-DNA was determined by protein induced fluorescence enhancement (PIFE) ^19^, where Cy3 fluorescence increases upon protein binding. Fluorescence spectra were acquired with a Fluoromax4 fluorometer (Horiba) using an excitation wavelength of 510 nm. 0.5 nM DNA was incubated for at least 5 minutes with 0-300 nM Reb1 in 10 mM Tris-HCl pH 8, 130 mM NaCl, 10% glycerol, 0.0075% v/v Tween-20. Fluorescence spectra were analyzed with Matlab to determine the change in Cy3 fluorescence.

### Ensemble FRET measurements

Reb1 binding to Cy3-Cy5 nucleosomes was measured as previously described ^4,42^. 0.5 nM nucleosomes were incubated for at least 5 minutes with 0-300 nM Reb1 in 10 mM Tris-HCl pH 8, 130 mM NaCl, 10% glycerol, 0.0075% v/v Tween-20. Fluorescence emission spectra were acquired as previously described ^42^. FRET efficiency was measured using the (Ratio)_A_ method ^43^.

### Single molecule TIRF microscope

The smTIRF microscope was built on an inverted IX73-inverted microscope (Olympus) as previously described ^20^. 532 and 638 nm diode lasers (Crystal Lasers) were used for Cy3 and Cy5 excitation. The excitation beams were expanded and then focused through a quartz prism (Melles Griot) at the surface of the quartz flow cell. A 1.3 N.A. silicone immersion objective (Olympus) was used to collect fluorescence, which were separately imaged onto an iXon3 EMCCD camera (Andor) with a custom built emission path containing bandpass filters and dichroic beam splitters (Chroma Tech). Each video was acquired using Micro-Manager software (Open Imaging).

### Flow Cell Preparation

Flow cells were functionalized as previously described ^44^. Briefly, quartz microscope slides (Alfa Aesar) were sonicated in toluene and then ethanol, and then further cleaned by piranha (3:1 mixture of concentrated sulfuric acid to 50% hydrogen peroxide). Slides were washed in water and, once completely dry, incubated in 100 uM mPEG-Si and biotin-PEG-Si (Laysan Bio) overnight in anhydrous toluene. Functionalized quartz slides and coverslips were assembled into microscope flow cells using parafilm with cut channels. Before each experiment, the flow cell is treated sequentially with 1 mg/ml BSA, 40 ug/ml streptavidin, and biotin-labeled DNA or nucleosomes.

### Single molecule fluorescence measurements of Reb1/Cbf1 binding kinetics

Biotinylated sample molecules (DNA or nucleosomes) were allowed to incubate in the flow cell at room temperature for 5 minutes and then washed out with imaging buffer containing the desired concentration of Reb1. The samples were first exposed to 638 nm excitation to determine the location of Cy5-labeled molecules and then 532 nm for both FRET and PIFE measurements. The imaging buffer for FRET experiments contained 10 mM Tris-HCl pH 8, 130 mM NaCl, 10% glycerol, 0.5% v/v Tween-20, 0.1 mg/ml BSA, 2 mM Trolox, 0.0115% v/v COT, 0.012% v/v NBA, 450 ug/ml glucose oxidase (Sigma G2133) and 22 ug/ml catalase (Sigma C3155), while the imaging buffer for PIFE experiments was contained 10 mM Tris-HCl pH 8, 130 mM NaCl, 10% glycerol, 0. 5% v/v Tween-20, 0.1 mg/ml BSA, 1% v/v BME, 450 ug/ml glucose oxidase (Sigma G2133) and 22 ug/ml catalase.

Single molecule time series were fit to a two-state step function by the hidden Markov method using vbFRET ^45^. Idealized time series were further analyzed using custom written Matlab programs to determine the dwell-time distributions of the TF bound and unbound states. 40% of traces were used in the analysis of FRET data and 13% of traces were used when analyzing PIFE data. Dwell-time and unbound-time cumulative sum distributions were generated from these traces and each distribution was analyzed using MEMLET to determine the best fit for the data and ultimately obtain rate constants for the transitions between bound and unbound states ^46^.

### FRAP Assay

Yeast cells containing GFP-labeled factors were cultured to log phase in synthetic medium, and then transferred to an agar pad and mounted by a coverslip. Cells were imaged using the 60× lens of FV1000 confocal microscope (Zeiss) at room temperature. 488nm laser was used to excite and bleach green fluorescence. Depending on the GFP intensity, 3-10% laser power was used to take images; 100% power was used to bleach the samples. The photobleaching time was set to 0.2 - 0.5s, and the intervals between consecutive frames were between 0.25 - 5s. Two frames were acquired before photobleaching, followed by 28-48 frames afterwards. The bleached region covered ~10%-25% of the nuclear region. Images were analyzed using Fiji-ImageJ. The average fluorescence intensity of bleached and unbleached regions was recorded, and the ratio between them was used in the recovery curve.

## Supplemental Figures

**S1:**
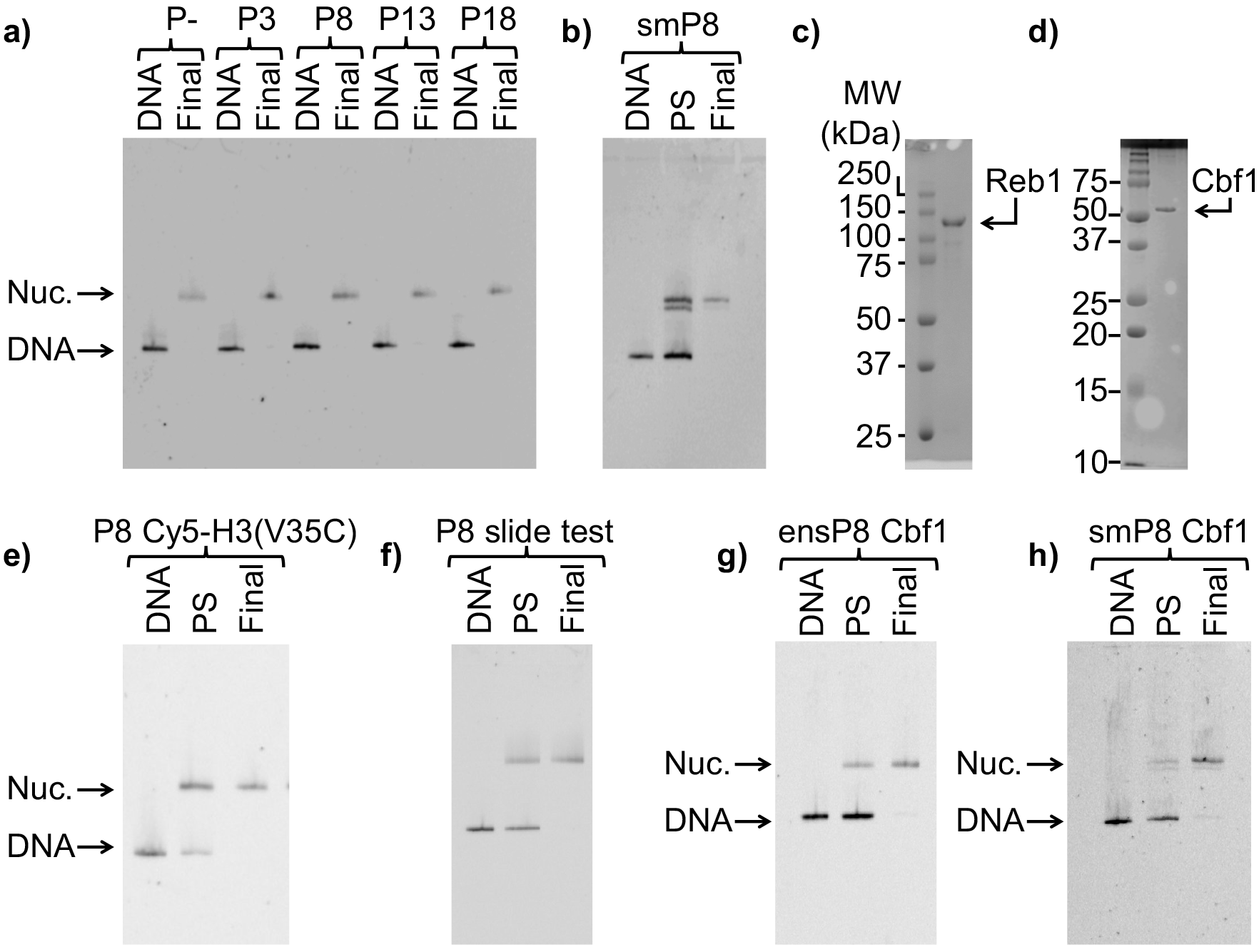
Nucleosomes and TFs used in this study. For all nucleosome gels: “DNA” = free DNA, “PS” = nucleosomes pre-sucrose gradient purification, and “Final” = final nucleosomes used for experiments, (**a**) Cy3 fluorescence image of the EMSA to quantify the purity of 147 bp reconstituted P-, P3, P8, P13 and P18 nucleosomes for ensemble Reb1-nucleosome experiments, (**b**) Cy3 fluorescence image of the EMSA of nucleosomes with the Reb1 binding site and a 75 bp linker for single molecule experiments, (**c**) Image of the the Coomassie stained of SDS-PAGE of purified Reb1 used in this study, (**d**) Image of the the Coomassie stained of SDS-PAGE of purified Cbf1 used in this study (**e**) Cy3 fluorescence image of the EMSA of the nucleosomes labeled with Cy5 at H3(V35C) for FRET, (**f**) Cy3 images of the EMSA of the nucleosomes used to test for Reb1-induced nucleosome repositioning. These nucleosomes contain a Reb1 binding site opposite the Cy3 fluorophore. We used 167 bp DNA for this assay, (**g**) Cy3 images of the EMSA of the 147 bp nucleosomes containing the Cbf1 binding site at P8 used for ensemble FRET experiments (**h**) Cy3 images of the EMSA of the 222 bp nucleosomes containing the Cbf1 binding site at P8 used for smFRET experiments.

**S2:**
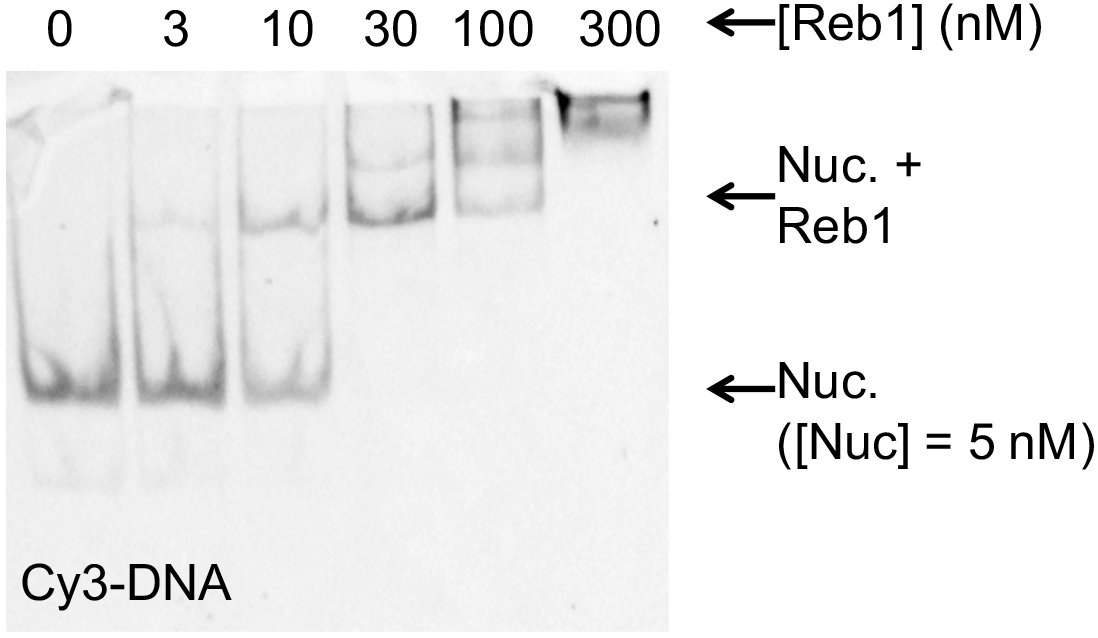
Rebl-nucleosome bound complex in EMSAs. Cy3 images of the EMSA of Reb1 binding to nucleosomes containing Cy5-H2A(K119C). This demonstrates the Reb1 shifted complex contains Cy3-DNA. This combined with Figure 1C demonstrates that the Reb1 binds a stable nucleosome complex.

**S3:**
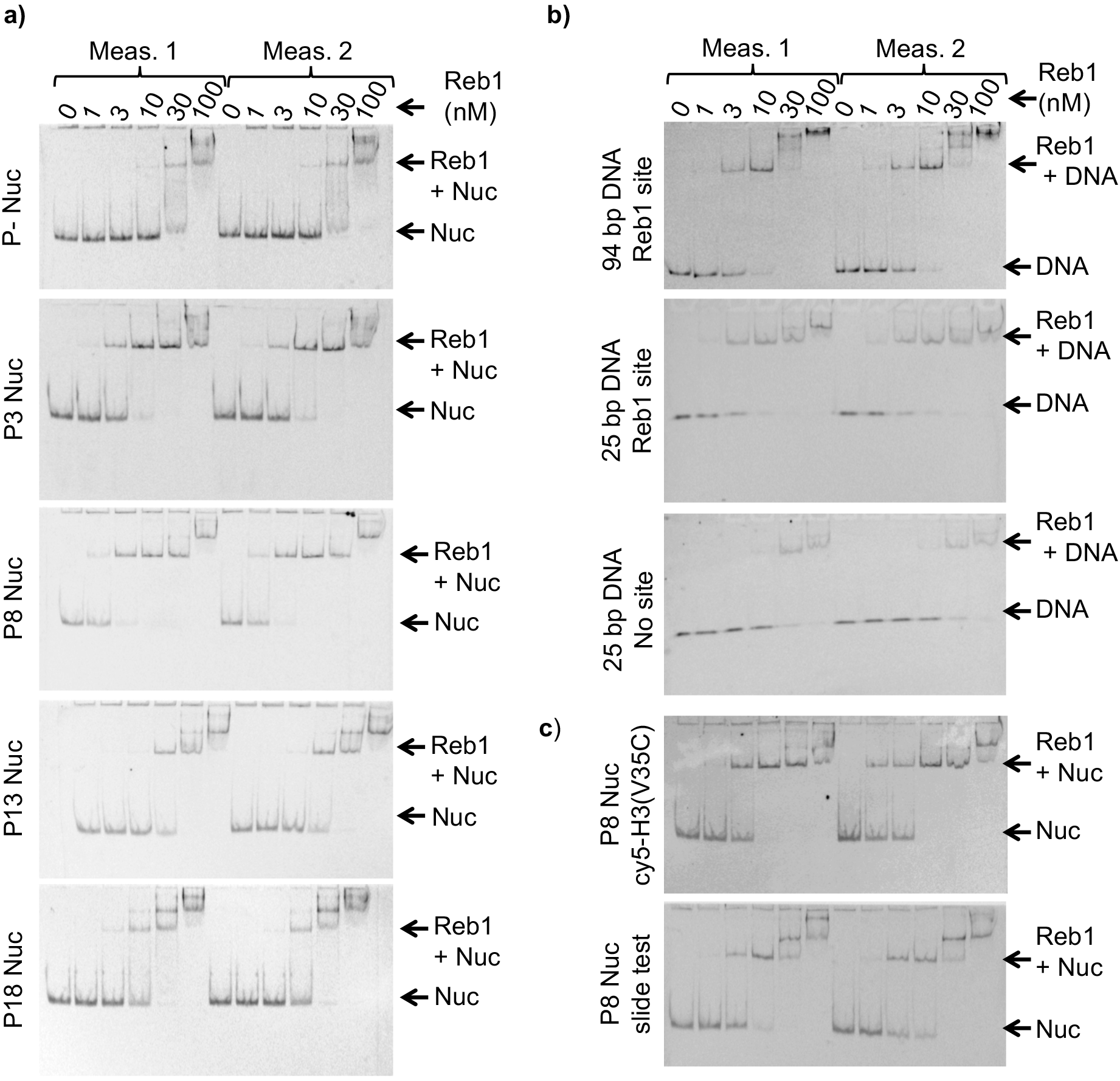
Nucleosome and DNA EMSAs. (**a**) Cy5 images of Reb1-nucleosome EMSAs. We define position of the binding site in the 601 sequence as“Px” where “x” represents the beginning of the Reb1 binding site. P- are nucleosomes without the Reb1 binding site. All EMSA measurements were performed in duplicate, (**b**) Cy3 images of EMSAs used to determine Reb1 binding to the DNA constructs with and without the Reb1 binding site. (**c**) Cy5 images of Reb1- nucleosome EMSAs with nucleosomes Cy5 labeled at H3(V35C) and Reb1-nucleosome EMSAs using nucleosomes to test for Reb1-induced nucleosome repositioning.

**S4:**
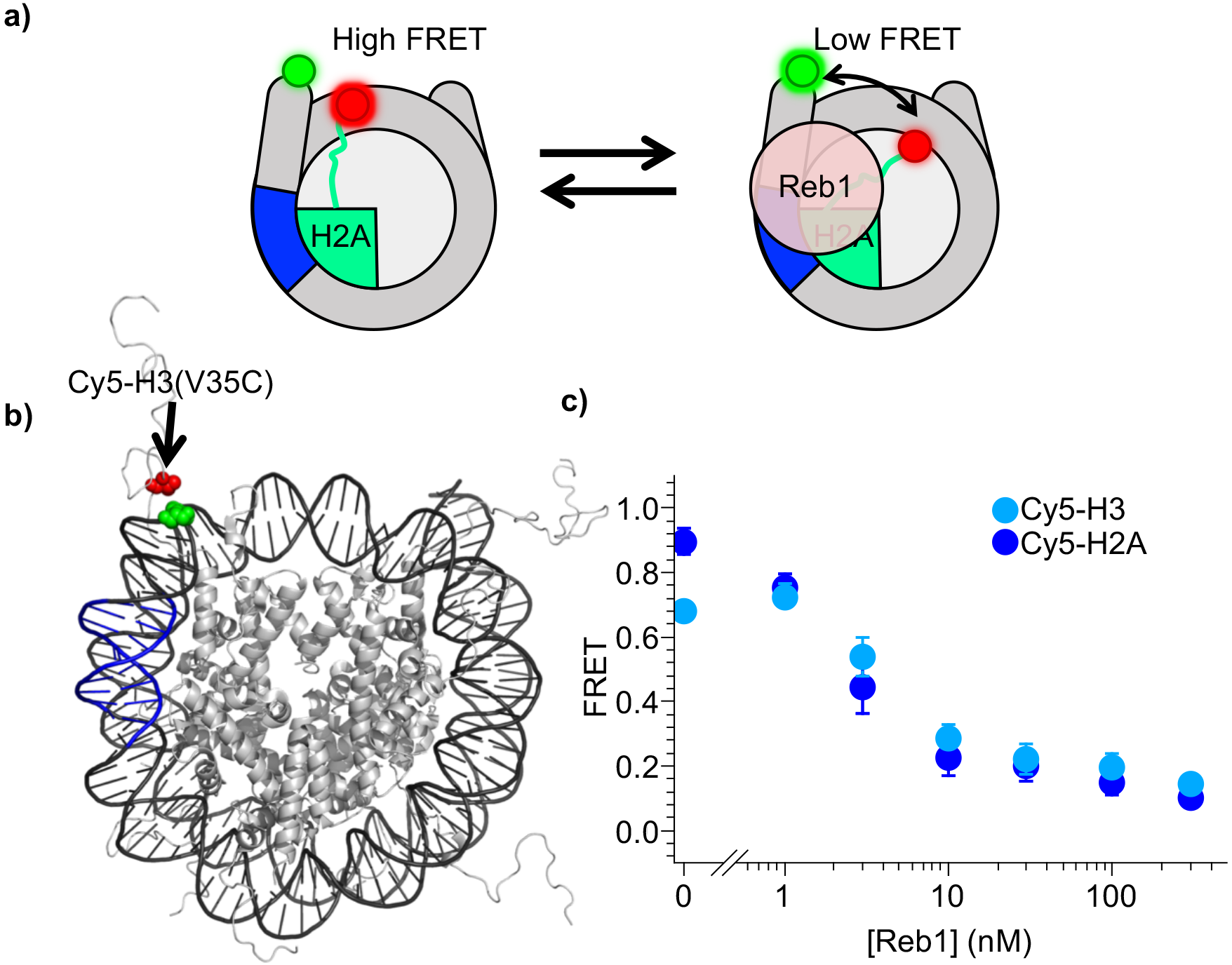
Reb1-induced nucleosome ΔFRET is not due to structural changes of the H2A C-terminal domain. (**a**) Nucleosome states representing one explanation for the observed Reb1-induced ΔFRET, which is that Reb1 distorts the H2A C-terminal domain, (**b**) Nucleosome structure ^38^ with the Reb1 binding site in blue, Cy3 in green and Cy5 that is attached to H3V35C shown in red. Cy3 and Cy5 undergoes efficient energy transfer when the nucleosome is fully wrapped, (**c**) Plot of the FRET efficiency of these nucleosomes with Reb1 titrated. We observe a similar decrease in FRET for Cy5-H3(V35C) (cyan) nucleosomes as with Cy5-H2A(K119C) (blue). This result demonstrates that the Reb1 dependent ΔFRET is not due to structural changes in H2A and instead is due to a structural change between the DNA and the entire octamer.

**S5:**
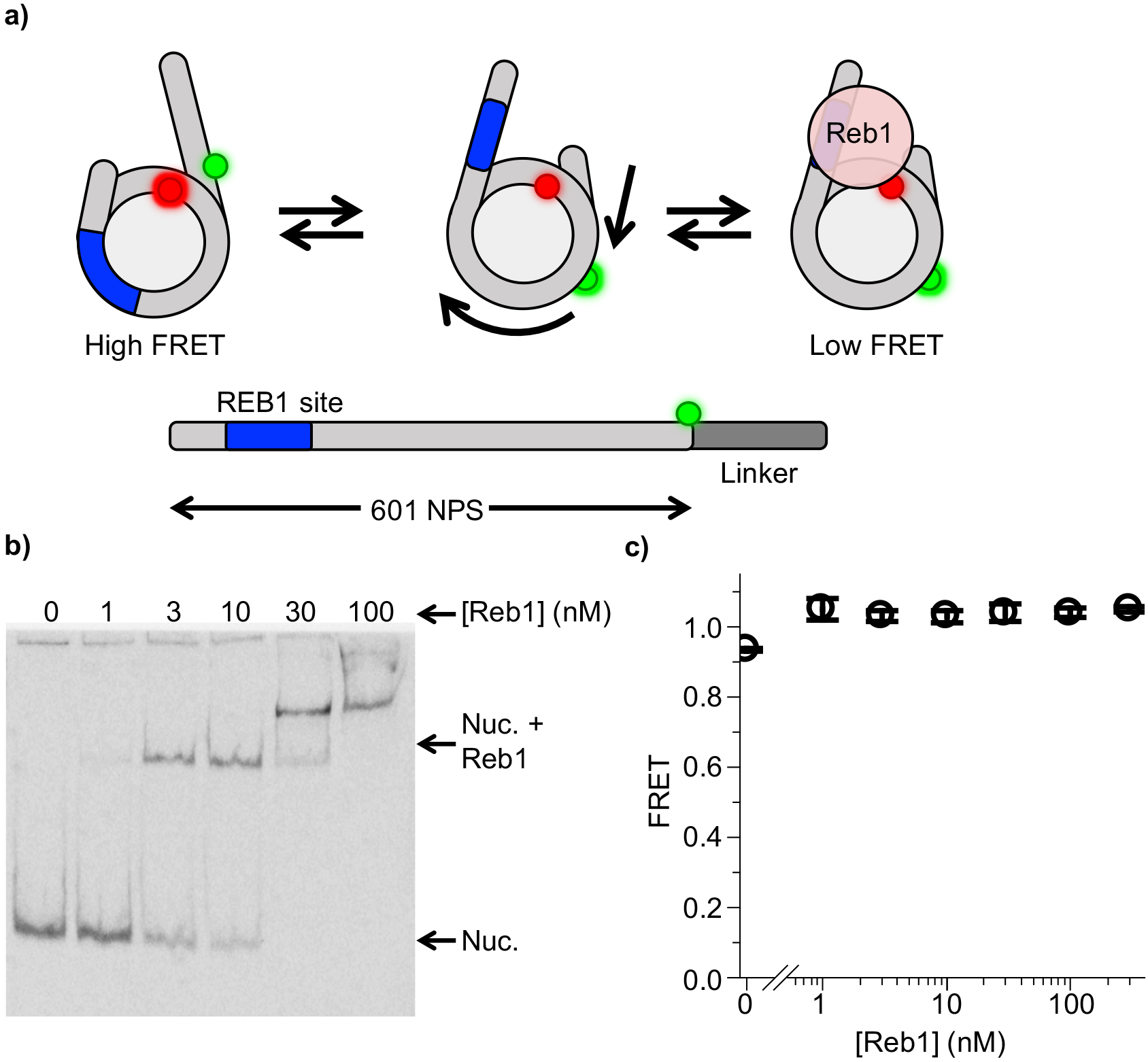
Reb1 does not trap nucleosomes in repositioned state. (**a**) Nucleosome states that represent a second possible explanation of the Reb1 dependent change in FRET, which is that Reb1 binds to repositioned nucleosomes that expose its binding site. To test this, we prepared 167 bp nucleosomes containing a Cy3 fluorophore and a 20 bp linker opposite the P8 Reb1 binding site. The octamer was labeled at H2A(K119C). (**b**) Cy3 image of the EMSA determining that Reb1 binds these nucleosomes. (**c**) A plot of the FRET efficiency from these nucleosomes as Reb1 is titrated. The FRET efficiency does not decrease as Reb1 is titrated over the concentration range that it binds nucleosomes. This result proves that the Reb1 dependent ΔFRET is not due to nucleosome repositioning.

**S6:**
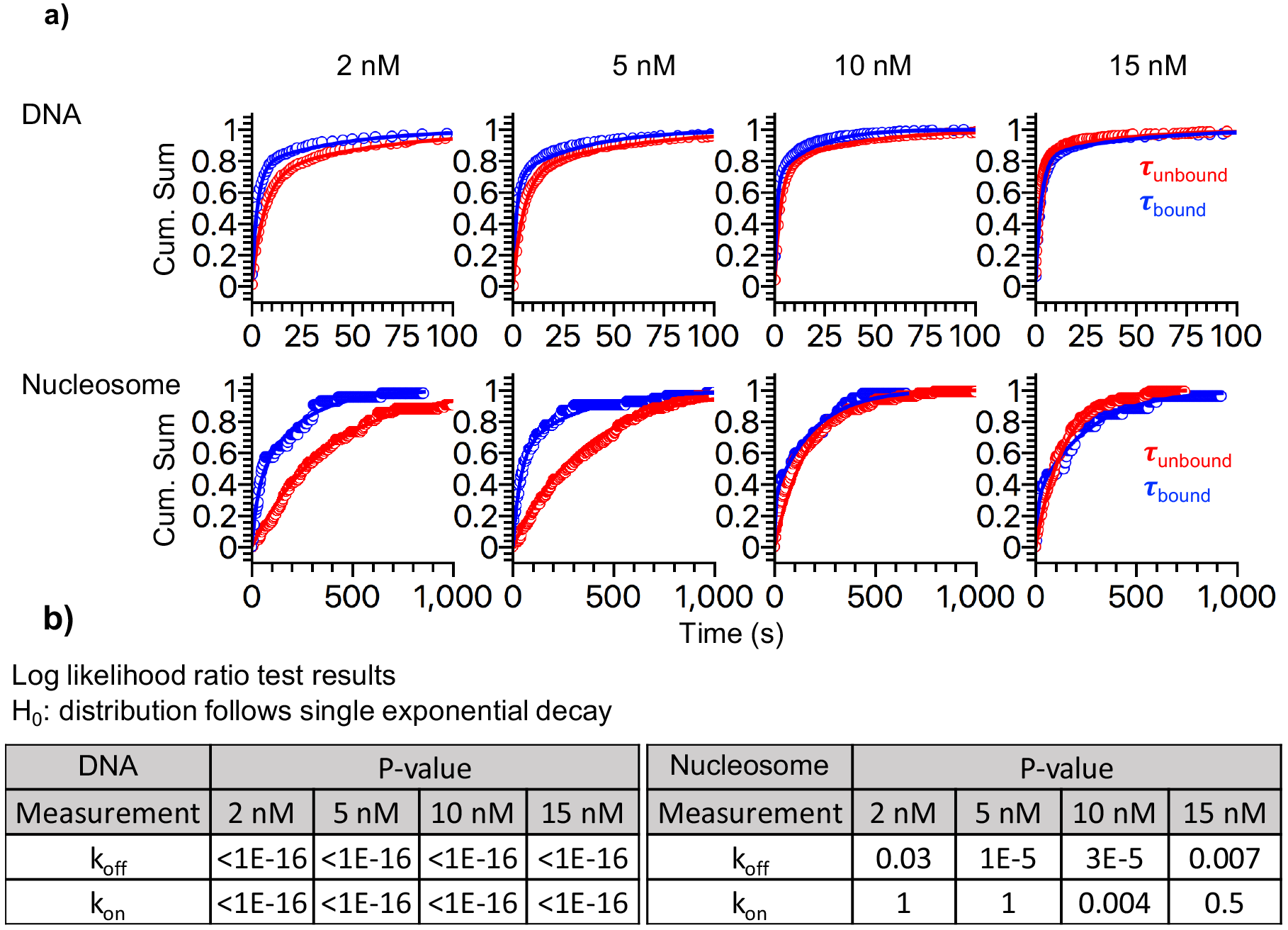
Reb1 binding and dissociation rates to DNA and nucleosomes. (**a**) Cumulative sums of unbound (red) and bound (blue) dwell times for DNA (top) and nucleosomes (bottom) for each Reb1 concentration, (**b**) Table of the log-likelihood ratio to test if the assumption that a cumulative sum fits to a double exponential. If the result is more than 0.01 for three of the Reb1 concentrations, we reject the assumption and fit with a single exponential. For Reb1-DNA, both unbound and bound cumulative sums follow double-exponentials. For Reb1-nucleosomes, the unbound cumulative sum fits best to a single exponential, while the bound cumulative sums fit best to a double exponential.

**S7:**
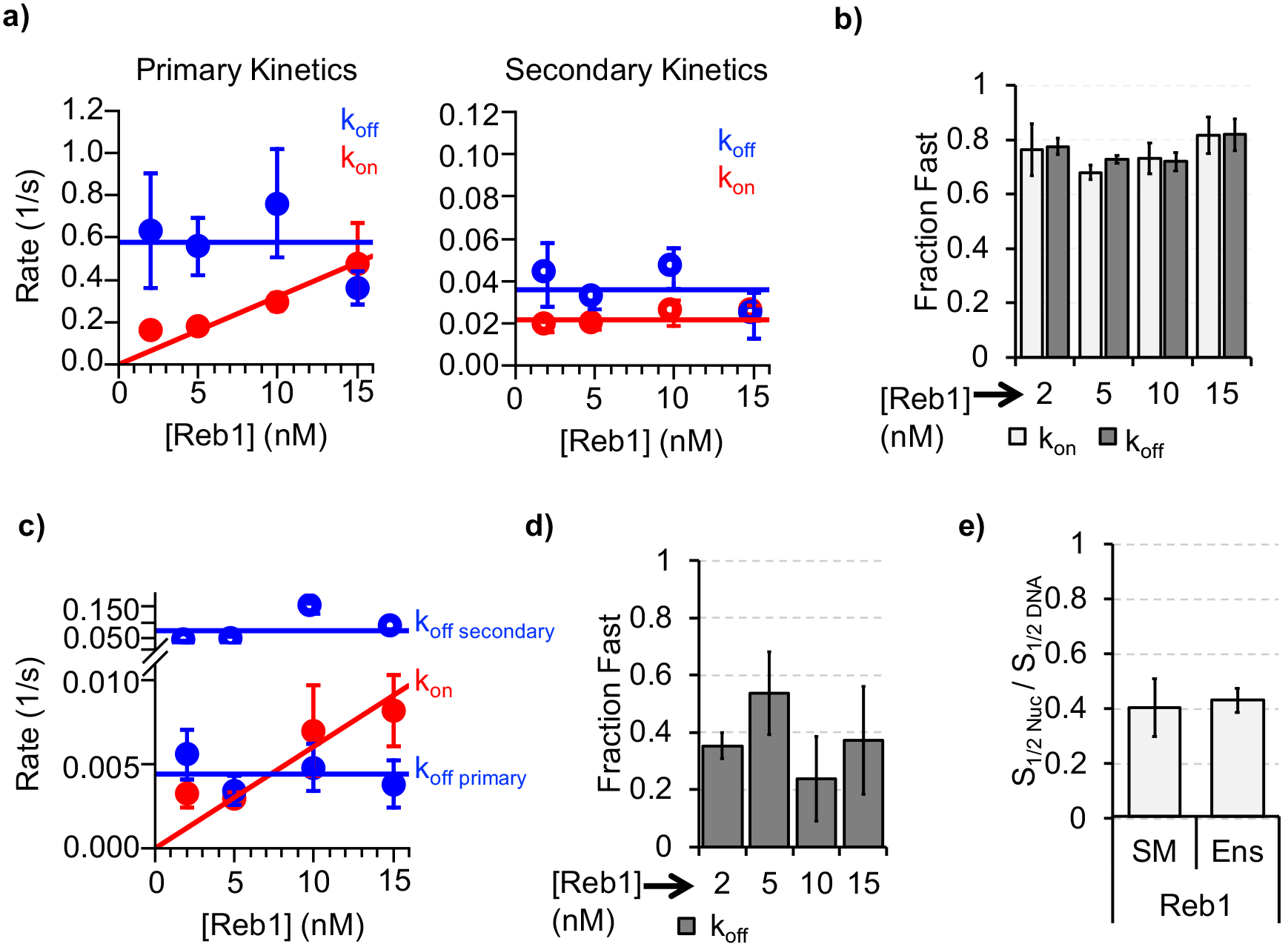
Analysis of Reb1-DNA and Reb1-nucleosome single molecule binding experiments. (**a**) Plots of primary and secondary rates for Reb1 binding and dissociation to and from DNA at increasing Reb1 concentrations. These are determined from the exponential fits of the cumulative sums in Figure S6. (**b**) The fraction of the fast binding (light grey) and fast dissociation rates (dark grey) of Reb1 to and from DNA for increasing Reb1 concentrations. For both Reb1 binding to and dissociating from DNA, the faster rates accounted for ~75% of the binding and dissociation events. (**c**) Plots of the Reb1 binding rates (red) to nucleosomes, and both the primary and secondary dissociation rates (blue) from nucleosomes for 4 concentrations of Reb1. These are determined from the exponential fits of the cumulative sums in Figure S6. (**d**) The fraction of the fast dissociation rates of Reb1 from DNA for increasing Reb1 concentrations. For Reb1 dissociation from nucleosomes, the slower rate accounted for ~60% of all of the dissociation events. (**E**) Ratio of nucleosome binding affinity to DNA binding affinity for Reb1 for both single molecule (SM) and ensemble (Ens) measurements. The ratio obtained using the dominant rates from single molecule measurements are consistent with the ratio from ensemble measurements.

**S8:**
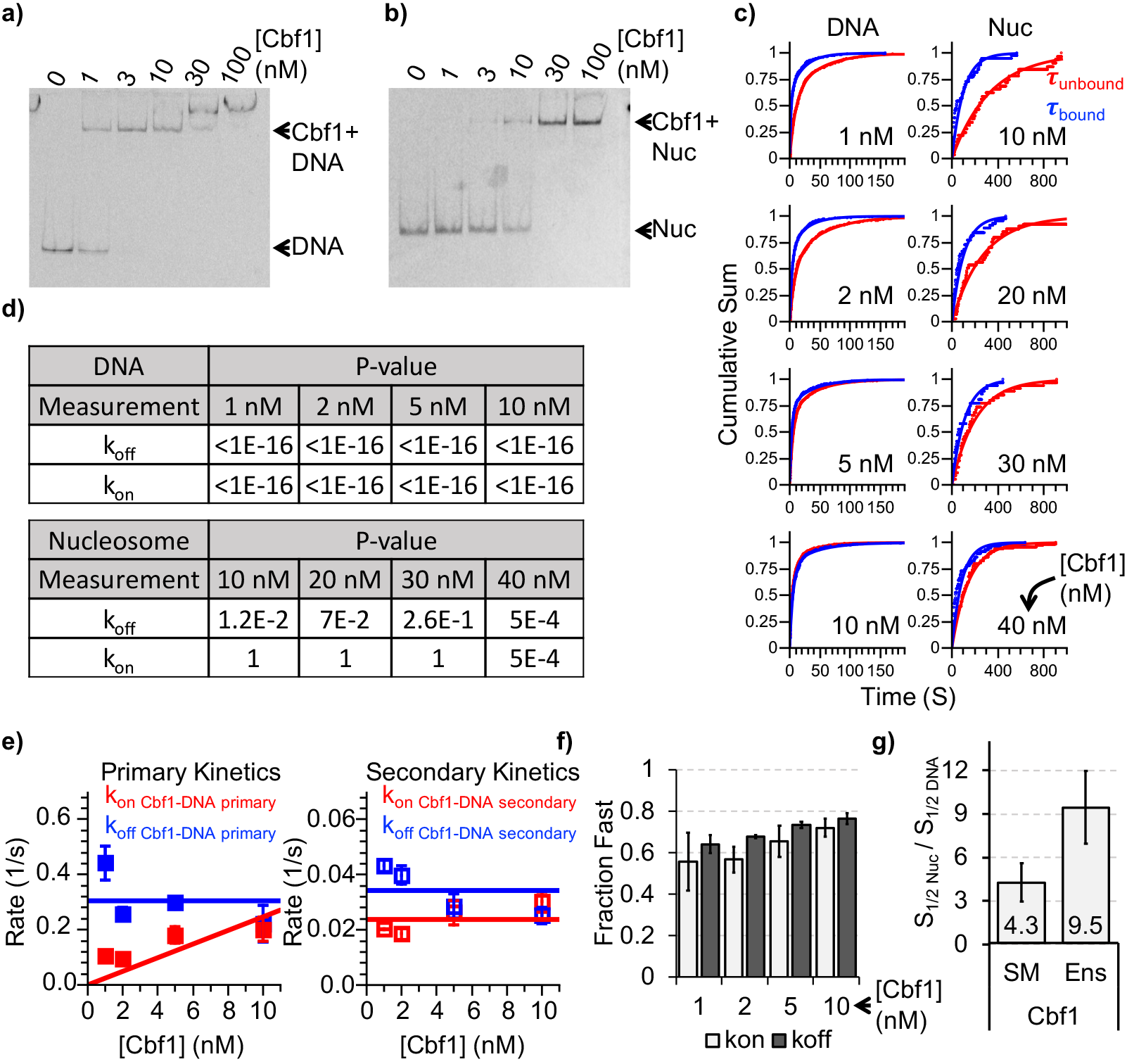
Characterizing Cbf1 interactions with DNA and nucleosomes. (**a**) Cy3 image of the EMSA of Cbf1 binding to a 94 bp DNA sequence with the Cbf1 binding site (S_1/2 cbfi-DNA EMSA_ ≈ 1 nM). (**b**) Cy5 image of the EMSA of Cbf1 binding to nucleosomes with the Reb1 binding site at P8 (S_1/2 cbfi-Nuc EMSA_ ≈ 11.5 nM). (**c**) Cumulative sum of unbound (red) and bound (blue) dwell times for DNA (left) and nucleosomes (right) for each Cbf1 concentration. The DNA cumulative sums were fit with double exponentials while the nucleosome cumulative sums were fit with single exponentials, (**d**) Table of the log-likelihood ratio to test if the assumption that a cumulative sum fits to double exponential. If the test is more than 0.01 for three of the four Reb1 concentrations, we reject the assumption and fit with a single exponential. For Cbf1-DNA, both unbound and bound cumulative sums follow double exponentials. For Cbf1-nucleosomes, both unbound and bound cumulative sums follow single exponentials, (**e**) Plots of primary (left) and secondary (right) rates for Cbf1 binding (red) and dissociation (blue) to and from DNA at increasing Cbf1 concentrations. These are determined from the exponential fits of the cumulative sums in Figure panel C. (**f**) The fraction of the fast binding (light grey) and fast dissociation rates (dark grey) of Cbf1 to and from DNA for increasing Cbf1 concentrations. For both Cbf1 binding to and dissociating from DNA, the faster rates accounted for ~60% and ~70% of the binding and dissociation events, respectively. (**g**) Ratio of nucleosome binding affinity to DNA binding affinity for Cbf1 for both single molecule (SM) and ensemble (Ens) measurements. The ratio obtained using the dominant rates from single molecule measurements is within about a factor of 2 of the ratio from ensemble measurements.

**S9:**
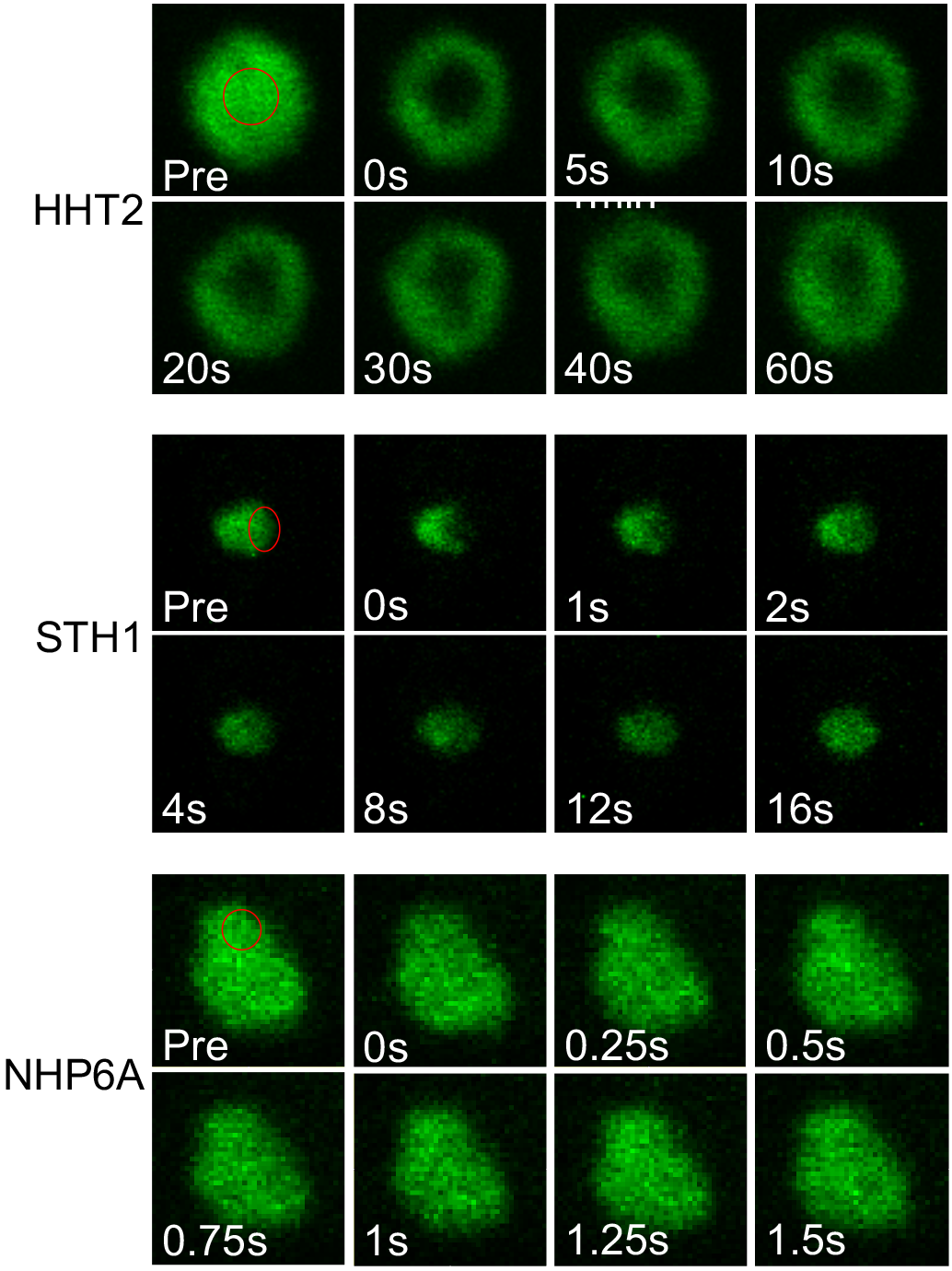
Fluorescence recovery of non-PF proteins in yeast nucleus. Fluorescence images of HHT2 (H3), STH1 and NHP6A during FRAP experiment. Bleached regions indicated with red circle.

**S10:**
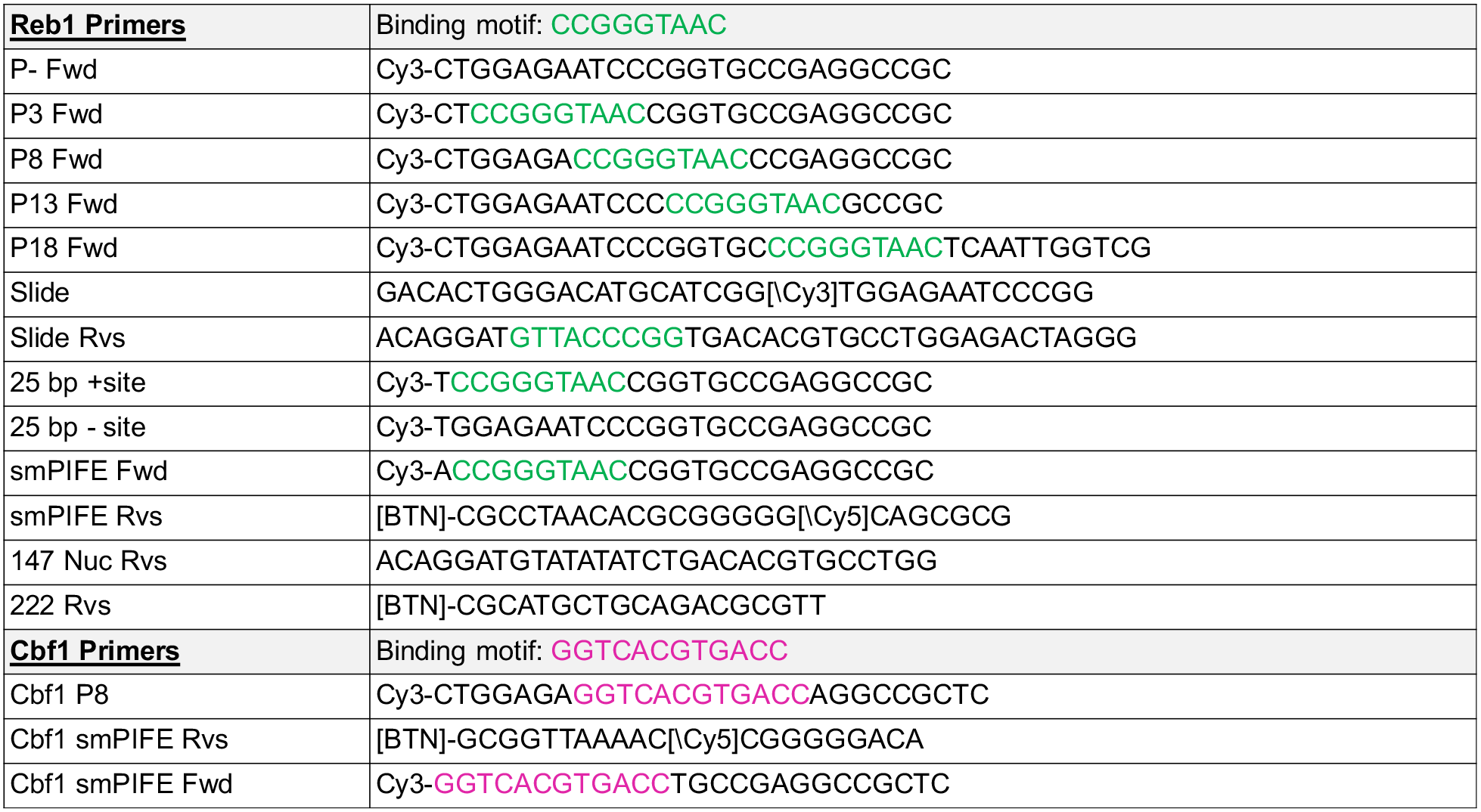
Oligonucleotides for preparing the DNA molecules used in this study.

